# Spatial activation of Kinesin-1 by Ensconsin shapes microtubule networks via ncMTOCs recruitment

**DOI:** 10.1101/2025.04.15.648882

**Authors:** Anne-Marie Berisha, Aude Pascal, Marine Guelle, Clément Bousquet, Denis Chrétien, Laetitia Bataillé, Régis Giet

**Author notes:** Lead contact: Régis Giet.

## Abstract

Intracellular transport, cell morphogenesis, motility, and division require the precise control of the microtubule (MT) cytoskeleton, which varies in shape and dynamic. *Drosophila* oogenesis requires the formation of cortical growing MTs that produces MTs twisters, sustaining cytoplasmic advection and polarization. Although Kinesin-1 MT motor is a central player regulating these processes, its exact function in the formation of these MTs is elusive. Here, we show that Ensconsin/MAP7, a Kinesin-1 activator, is transported by Dynein together with MTs and maintained at the oocyte cortex by Ninein to regulate spatial activation of Kinesin-1 in the oocyte. Perturbation of this process leads to a severe reduction of ncMTOCs (non-centrosomal MicroTubule Organizing Centers) targeting to the cell cortex, and reduced MTs anchoring and stabilization. We also show that an active Khc variant, unable to bind Ens/MAP7 and harboring a loss of auto-inhibition, is sufficient to restore ncMTOCs recruitment, MTs twister formation and advection in the oocyte. Our findings reveal a pivotal mechanism by which the targeted localization of activators drives the spatial activation of motors, a process fundamental to microtubule remodeling.

## Introduction

Multicellular organisms possess a remarkable ability to reorganize their cytoskeleton, enabling the formation of specialized cells with distinct shapes and functions. Among these, the production of functional gametes stands out as a prime example of the intricate microtubule (MT) rearrangements required. In *Drosophila,* oogenesis begins with four divisions of a single stem cell, producing a 16-cell cyst interconnected by ring canals resulting from incomplete cytokinesis. A single cell forms the future oocyte, while the other 15 cells become polyploid, act as the oocyte’s nurse cells (NCs) that provides nutrients, proteins, mRNA and organelles.

These components are transported through ring canals to ensure the oocyte growth (Robinson and Cooley, 1996). Studies have shown that dynein’s minus-end directed motor activity, together with that of the actin binding protein Short Stop (Shot) at ring canals, can orient and allow the passage of MTs from NC to the oocyte, contributing to a cytoplasm and organelle flow that sustains oocyte growth (Lu et al., 2022).

In parallel of the growth process, the polarization of the oocyte is achieved *via* the transport and anchoring of mRNA to specific locations during the mid-oogenesis stages (Parton et al., 2011; Sanghavi et al., 2016). For example, the polarization of *oskar* mRNA at the posterior site of the oocyte determines the antero-posterior axis of the future embryo. This transport is MT dependent and involves the plus-end directed Kinesin-1 motor activity (Brendza et al., 2000; Brendza et al., 2002; Serbus et al., 2005). It has long been thoughted that Khc (Kinesin-1 heavy chain)-mediated transport of mRNAs toward posterior cortex occurs on a polarized cortical MT network. Indeed, examination of MT plus ends growth have shown that MTs exhibit a low but sufficient bias in MT orientation towards the posterior cortex of the oocyte (Parton et al., 2011; Zimyanin et al., 2008). How this cortical MT network is established in the oocyte is still not well understood. MT nucleation at the oocyte cortex does not require the conventional γ-tubulin, that is associated with canonical MTOC (MicroTubule Organizing Centers), such as centrosomes. Instead, MT organization during mid-oogenesis relies on ncMTOCs (non-centrosomal MicroTubule Organizing Centers). ncMTOCs are composed of the MT minus end binding protein Patronin which is associated with the spectraplakin protein Shot, anchored to the cortical actin cytoskeleton (Nashchekin et al., 2016; Quinlan, 2016; Tillery et al., 2018).

At the transition between mid-oogenesis to late oogenesis, a cortical MT network of long and parallel MTs appears taking the form of streams (or twisters) that stir the cytoplasm to create advection and a homogeneous distribution of organelles in the oocytes, and contributes to complete *oskar* mRNA anchoring at the posterior cortex (Lu et al., 2018). How this collective movement of ooplasm is established by thousands of MTs has been recently modelled: Kinesin-1-dependent cargo transport on a single MT is able to move surrounding cytoplasm and therefore neighboring MTs. MTs then become collectively bent to create MT streams or twisters, and Kinesin-1 motor transport of cargoes toward MT plus ends generate a robust flux of cytoplasmic components in the oocyte (Dutta et al., 2024; Monteith et al., 2016). Supporting this model, lowering Khc activity generates a slower flux: ooplasmic advection is therefore a solid readout of Khc activity (Ganguly et al., 2012). However, Khc may controls other mechanisms leading to cortical MT arrays formation.

How Kinesin-1 is specifically activated in the oocyte remains a crucial but elusive question. A previous study demonstrated that Khc is evenly expressed in both nurse cells (NCs) and oocytes but has no functional role in NCs (Brendza et al., 2000; Brendza et al., 2002), strongly suggesting that the spatial activation of Kinesin-1 is not dependent on Khc levels. A key regulatory mechanism responsible for this spatial activation of Khc may involve the MAP7 (Microtubule-Associated Protein 7) family member Ensconsin (Ens) (Barlan et al., 2013; Hooikaas et al., 2019; Metivier et al., 2019; Tymanskyj et al., 2018). Indeed, the MAP7 family of proteins has been shown to recruit Khc to microtubules (Hooikaas et al., 2019; Metivier et al., 2019; Monroy et al., 2018; Sung et al., 2008). Furthermore, Ens was initially reported to be enriched in the oocyte, positioning it as a strong candidate for mediating the spatial activation of Khc (Sung et al., 2008).

Here we dissected the mechanisms that lead to the spatial enrichment of Ensconsin and showed that this enrichment controls Khc activation in the oocyte. We also show that this spatial activation of Khc is required for ncMTOCs targeting to the cell cortex, a founding event that sustains cortical MT twisters’ assembly. This result demonstrates the role of the Khc/Ens as not only the main user but also a builder of its own cortical MT network.

## Results

### Dynein promotes the enrichment of the Khc activator Ensconsin in the oocyte

The egg chamber consists of interconnected cells where Dynein and Khc activities are spatially separated. Dynein facilitates transport from NCs to the oocyte, while Khc mediates both transport and advection within the oocyte. To identify the mechanisms regulating Khc activation in the oocyte, we first analyzed its spatial distribution within the egg chamber. Immunostaining with anti-Khc antibodies confirmed that Khc is present in both NCs and the oocyte (Figure S1A), consistent with previous findings (Brendza et al., 2000; Brendza et al., 2002). Similarly, a functional Khc^WT^-GFP expressed at similar levels as endogenous Khc displayed the same distribution between NCs and the oocyte (Figure S1B, and Figures 1A top left and 1B). We therefore confirm that Kinesin-1 activity in the oocyte is not driven by Khc spatial enrichment. A previous studies have revealed that the recruitment and activity of Khc on MTs are governed by the binding to MAP7 family of proteins (Barlan et al., 2013; Hooikaas et al., 2019; Metivier et al., 2019; Tymanskyj et al., 2018). Thus, we analyzed the localization of Ensconsin, the sole MAP7 family member present in the *Drosophila* genome. A functional Ens^WT^-GFP transgenic protein displayed a strong enrichment in the oocyte, in agreement with early observations (Figures 1A top right and 1B, (Metivier et al., 2019; Sung et al., 2008)).

Therefore, we analyzed the molecular mechanisms by which Ens is enriched in the oocyte. A previous study showed that the 3’ UTR of the *ens* mRN A displays an oocyte localization signal but this sequence is dispensable for oocyte protein enrichment (Sung et al., 2008). This suggests that the Ensconsin protein itself harbors motifs that mediate this enrichment. Ens display two conserved functional domains: a C-terminal domain able to bind Khc and a N-terminal domain that mediates MTs binding (Figure S1C, (Barlan et al., 2013; Hooikaas et al., 2019; Metivier et al., 2019)). We generated and expressed specific Ens variants to quantify the impact of the MT and Khc-binding properties for oocyte enrichment (Figures S1C and S1D). An Ens^MutKhc^ with lower affinity for Kinesin-1 was generated by mutating a putative Khc binding motif in the C-terminal domain of Ens (Figure S1C, (Hooikaas et al., 2019; Monroy et al., 2018)). Unlike Ens^WT^, the Ens^MutKhc^ variant did not interact with full-length Khc (Figure S1E). When expressed in the oocyte, Ens^MutKhc-GFP^ showed an enrichment pattern similar to that of its WT counterpart, suggesting that Ens targeting in the oocyte is independent of Khc (Figures 1A, bottom left, and 1B). Consistently, Khc depletion by RNAi did not abolish Ens enrichment in the oocyte (Figures S1F and S1G).

**Figure 1:**
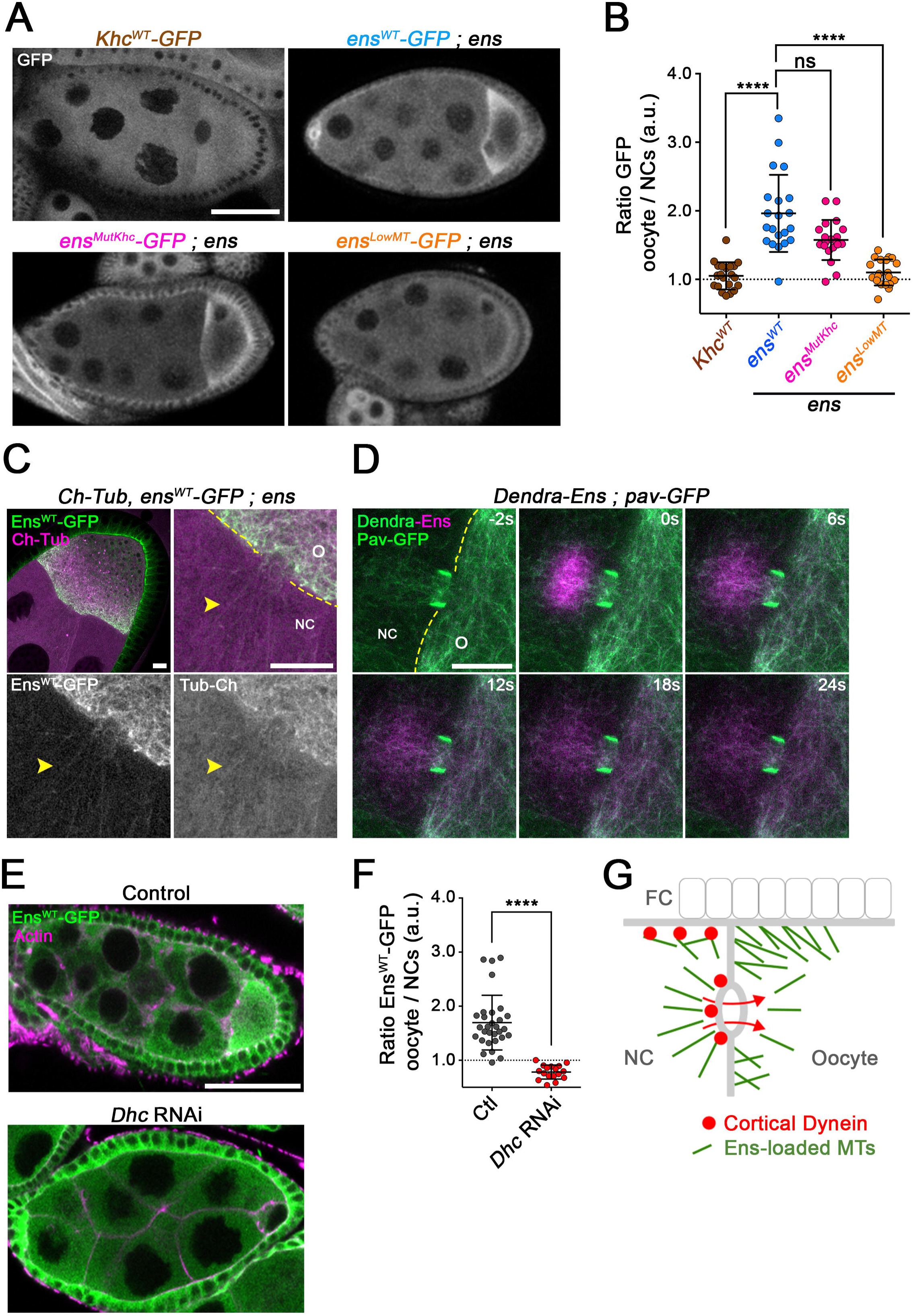
Mechanism of Ens enrichment in the oocyte. **(A)** Images of Khc^WT^-GFP (top, left) and *ens* mutant egg chambers expressing either Ens^WT^-GFP (top, right), Ens^MutKhc^-GFP (bottom, left), or Ens^LowMT^-GFP (bottom, right), showing the GFP distribution in nurse cells and oocytes during stage 8. Images are 3 µm projections. Scale bar: 50 µm. **(B)** Scatter dot plot showing the mean GFP intensity ratio (± sd) between the oocyte and NC in egg chambers expressing Khc^WT^-GFP (1.05 ± 0.20, *n =* 21), or in *ens* mutant expressing Ens^WT^-GFP (1.96 ± 0.56, *n =* 21), Ens^MutKhc^-GFP (1.57 ± 0.29, *n =* 20) and Ens^LowMT^-GFP (1.10 ± 0.19, *n* = 20). ns, not significant; ****: p<0.000l (Kruskal-Wallis non-parametric test and Dunn’s multiple comparisons tests). **(C)** Stage 9 *ens egg* chamber expressing Ch-Tubulin and Ens^WT^-GFP. Ens^WT^-GFP (green in the merge panels) and Ch-Tubulin (magenta in the merge panels) are also shown in monochrome in the bottom panels. The yellow arrowheads point to Ens-GFP decorated MTs in the NC. Here and below, the yellow lines indicate the boundary between the NC and the oocyte (O).. Scale bars: 10 µm. **(D)** Selected time frames from a photoconversion experiment on a stage 9 egg chamber expressing Dendra-Ens and Pav-GFP. Pav-GFP is shown in green and labels the ring canal. Time t = 0s is the photo-conversion starting point and shows the pool of the photoconverted Dendra-Ens MTs (magenta) that reach the oocyte trough the time course of the experiment. Time in s is indicated for each panel. Scale bar: 10 µm. See also supplementary Video 1. **(E)** Ens^WT^-GFP distribution in control (top) and *Dhc* RNAi (bottom) egg chambers, stained for GFP (green) and actin (magenta). Ens^WT^-GFP accumulation in the oocyte is compromised following depletion of Dhc. Scale bar: 50 µm **(F)** Scatter dot plot showing the mean Ens^WT^-GFP intensity ratio (± sd) between the oocyte and NC in control (1.70 ± 0.51, *n =* 30) and *Dhc* RNAi (0.78 ± 0.13, *n =* 17) conditions. ****: p<0.000l (Mann-Whitney test). **(G)** Schematic representation of Ens-loaded microtubules travelling from a nurse cell toward the oocyte via the ring canal thanks to Dynein. NC: nurse cell and FC: follicle cells.

In parallel, to investigate if Ens MT-binding properties may influence its localization, we generated an Ens variant with lower affinity for MTs. To achieve this, we took advantage of previous observations showing that Human Ensconsin (MAP7) binding to MT is down regulated following phosphorylation of the MT-binding domain by the mitotic kinase Cdk1 (Bulinski et al., 2001; McHedlishvili et al., 2018). We thus identified the consensus Cdk1 phosphorylation sites on Ens MT-binding domain and generated phospho-mimicking mutations on Ens to generate an Ens^LowMT^ variant (Figure S1C). When compared to the WT variant, this Ens^LowMT^-GFP variant harbored a significantly loss of localization on MTs when expressed in oocytes (Figures S1H and S1I) and a decreased ability to polymerize MTs in turbidity assays (Figure S1J). In egg chamber, this Ens^LowMT^-GFP displayed a significant decreased oocyte enrichment compared to Ens^WT^-GFP indicating that Ens binding to MTs controls its spatial distribution (Figure 1A bottom right and 1B). Consistent with this observation, Ens-GFP co-localized with the MTs in NC, as well as with those crossing the ring canals between NC and the oocyte (Figure 1C, yellow arrowhead). Moreover, when photo-converted in NCs, in a region adjacent to the ring canals, Dendra-Ens-labeled MTs were found in the oocyte compartment indicating they are able to travel from NC toward the oocyte trough ring canals (Figure 1D and Video 1). Transport of MTs from NCs toward the oocyte trough ring canals is mediated by Dynein (Lu et al., 2020; Lu et al., 2022). Dynein heavy chain (Dhc) RNAi completely abolished enrichment of Ens-GFP in the oocyte as well as oocyte growth (Figures 1E and 1F).

Altogether, these data support a mechanism by which Ens loading on MTs in NCs and their subsequent transport by Dynein toward ring canals promotes the spatial enrichment of the Khc activator Ens in the oocyte (Figure 1G).

### Ninein mediates Ens localization in the oocyte

To further characterize the mechanisms of Ens enrichment, we also examined the possible contribution of the *Drosophila* Ninein (Nin, also known as Bsg25D) to this process. This MAP has been described to physically bind Ens and regulate its localization and function in differentiated cells (Rosen et al., 2019; Tillery et al., 2024). We therefore analyzed Ninein localization during oogenesis. At stage 9, Ninein was partially co-localized with Ens-associated MTs at the latero-anterior cortex (Figure 2A), a position where the ncMTOC component Shot resides to organize cortical MTs (Nashchekin et al., 2016). Then at stage 10, a step in which cortical MT streams are generating cytoplasm advection, Ninein was barely detected in association with any structure within the oocyte (not shown, (Kowanda et al., 2016)).

**Figure 2:**
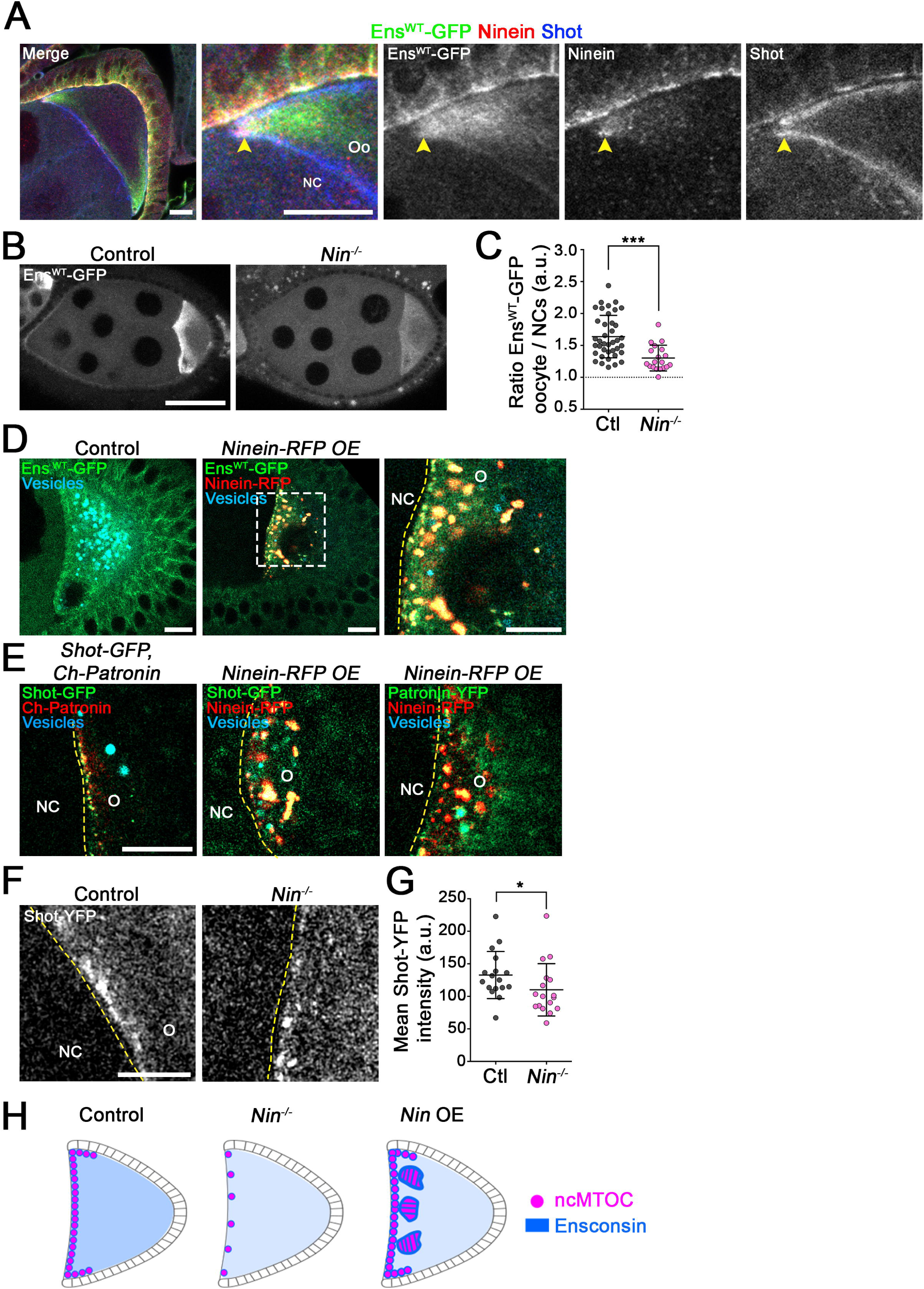
Ninein promotes Ens enrichment and favors ncMTOCs cortical recruitment in the oocyte. **(A)** Stage 9 Ens^WT^-GFP oocyte stained for GFP (green in the merge channel), Ninein (red) and the ncMTOC component Shot (blue). Indicated proteins are also shown in monochrome. Scale bars: 10 µm. The arrow head points to the cortical region where the three proteins co-localize. **(B)** Ens^WT^-GFP distribution in a control (left) and a *Ninein* mutant (right) egg chamber. Scale bar: 50 µm. Loss of Ninein leads to a decreased enrichment of Ens^WT^-GFP in the oocyte. **(C)** Scatter dot plot showing the mean Ens^WT^-GFP intensity ratio (± sd) between the oocyte and NC in control (1.64 ± 0.33, *n =* 39) and in *Ninein* mutant stage 8 egg chambers (1.3 ± 0.20, *n = 19). ***: p<0.00l* (Unpaired t-test). **(D)** Ens^WT^-GFP distribution in control (left) and in Ninein-RFP overexpressing stage 9 oocytes (middle). The white dotted square shows the region shown in the right panel. Increased levels of Ens^WT^-GFP (green) were recruited onto Nin-RFP aggregates (red) at the cortex (n = 14 oocytes). Yolk vesicle autofluorescence are shown in cyan. Scale bars: 10 µm. **(E)** Localization of the ncMTOC components Shot and Patronin following Ninein overexpression. In a control oocyte (left) Shot-GFP (green) and Ch-Patronin are located at the anterior cortex. In Ninein-RFP overexpressing oocytes (middle and right panels), Shot-GFP (green, middle panel) and Patronin-YFP (green, right panel) are recruited to Nin-RFP aggregates at the cortex. Yolk vesicles autofluorescence are shown in cyan. The recruitment of ncMTOC proteins on Ninein cortical aggregates was observed in 100 % of the oocytes (n>11 oocytes). Scale bar: 10 µm. **(F)** Cortical anterior localization of Shot-YFP in control (left) or *Ninein* mutant (right) stage 9 oocytes. Loss of Ninein leads to loss of Shot-YFP at the anterior cortex. Scale bar: 10 µm. **(G)** Scatter dot plot showing the mean Shot-YFP intensity (± sd) at the anterior cortex, in control and in *Ninein* mutant stage 9 oocytes. Control (132.7 ± 36.22, *n =* 17), *Ninein* mutant (110.1 ± 40.22, *n =* 17). *: p<0.05 (Mann-Whitney test). **(H)** Schematic representation of ncMTOCs (magenta) distribution in a control (left), a *Ninein* mutant (middle) and a Ninein overexpressing stage 9 oocyte (right). The blue background color reflects Ens levels in the oocyte.

Given the physical interaction between Ens and Ninein and their partial colocalization in the oocyte, we investigated if Ensconsin localization was affected by the loss of Ninein. In agreement with previous studies, *Ninein* mutant females were viable and fertile (Kowanda et al., 2016; Rosen et al., 2019; Tillery et al., 2024). *Nin* oocytes harbored a WT growth, indicative of a functional dynein transport and consequently a normal oocyte feeding trough the ring canals. However, Ensconsin enrichment was strongly diminished in these oocytes (Figures 2B, 2C and 2H). In a reciprocal test, to assay if elevation of Ninein levels was able to mis-regulate Ens localization, we overexpressed a tagged Ninein-RFP protein in the oocyte. At stage 9 the overexpressed Ninein accumulated at the anterior cortex of the oocyte and also generated large cortical aggregates able to recruit high levels of Ens (Figures 2D and 2H). These aggregates were not maintained over time as they were barely detected at stage 10 and did not affect female fertility (Figures S2A and Table S1).

In conclusion, Ninein co-localizes with Ens at the oocyte cortex and partially along cortical microtubules, contributing to the maintenance of high Ens protein levels in the oocyte and its proper cortical targeting.

### Nin contributes to ncMTOC recruitment to the cell cortex

During mid-oogenesis (stage 9), the recruitment of the ncMTOC proteins Shot and Patronin at the antero-lateral cortex, is required to anchor MTs by their minus ends and mediate MT organization (Nashchekin et al., 2016). The co-localization of Ninein together with Ens, and Shot at the oocyte cortex (Figure 2A), prompted us to monitor a possible effect of Ninein on the dynamics of Shot and Patronin recruitment. The examination of Ninein/Ens cortical aggregates obtained after Ninein overexpression showed that these aggregates were also able to recruit high levels of Patronin and Shot (Figures 2E and 2H). Reciprocally, analysis of cortical ncMTOCs recruitment revealed that *Nin* mutant oocytes displayed a significant loss of cortical Shot in stage 9 oocytes (Figures 2F, 2G and 2H). To investigate if this lower Shot recruitment during early oogenesis could impact MT organization at later stages, we monitored the presence of MT streams (Figure S2B) and quantified cytoplasmic advection that occurs during late oogenesis at stage 10B, as this is a direct readout of anchored MT twister assembly (Figure S2C, (Ganguly et al., 2012)). We found that only 50% of *Nin* oocytes displayed the presence of cortical MT streams (Figure S2B) that were associated with a low advection. Taken together, these results indicate that, although non-essential, Ninein contributes to recruit Ens and ncMTOCs at the oocyte cell cortex during mid-oogenesis and contributes to MT-dependent streaming at later stages.

### Ens and Khc are essential for ncMTOC cortical localization

Given our findings that Ninein is required to maintain high Ensconsin levels in the oocyte and to recruit cortical ncMTOCs, we tested whether Ensconsin and Khc could mediate ncMTOC cortical formation. Observation of lives stage 9 oocytes, revealed that Ens-GFP was strongly associated with the cortical InfraRed-labelled MTs (Figure 3A). Moreover, a pool of the Ens-GFP co-localized with Ch-Patronin at cortical ncMTOCs at the anterior cortex (Figure 3A). The presence of Ens with Patronin and Shot at ncMTOCs (Figure 2A and Figure 3A) prompted us to examine if these proteins were physically associated. Co-immunoprecipitations experiments revealed that Patronin was associated with Shot-YFP, as shown previously (Nashchekin et al., 2016), but also with Ens^WT-^ GFP, indicating that Ens, Shot and Patronin are present in the same complex (Figure 3B).

**Figure 3:**
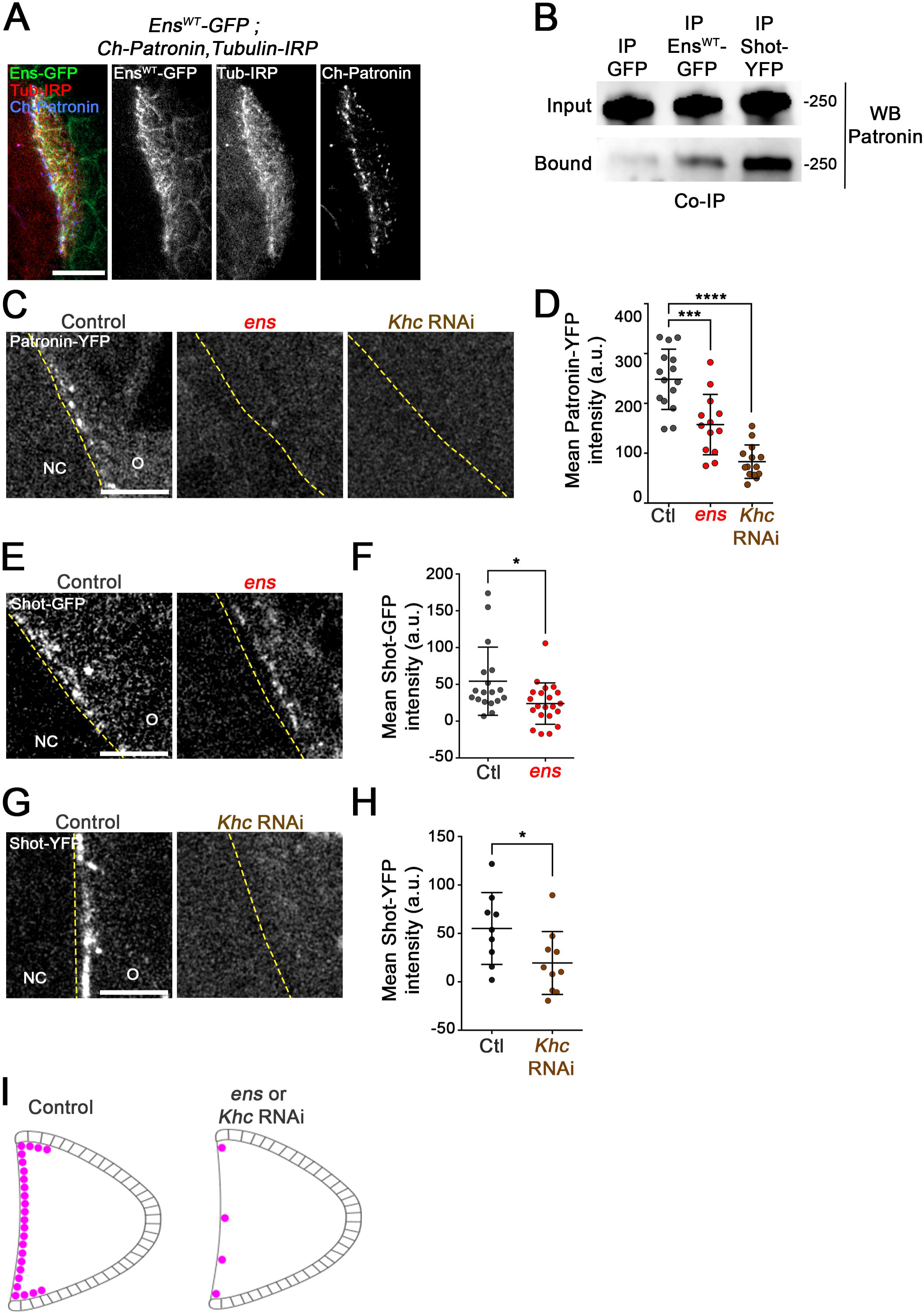
Ens and Khc are required for ncMTOC cortical recruitment. **(A)** Localization of Ens^WT^-GFP (green), Tubulin-InfraRed (red) and Ch-Patronin (blue) in a stage 9 oocyte, showing the localization of Ens along MTs, and its colocalization with Patronin at the anterior cortical MTs anchoring sites. **(B)** Western blot analysis of GFP, Ens^WT^-GFP, and Shot-YFP immunoprecipitates using an anti-Patronin antibody. Patronin is detected in Ens^WT^-GFP and Shot-YFP but not GFP immunoprecipitates. **(C)** Patronin-YFP cortical localization in a control (left), *ens* mutant (middle) and *Khc* RNAi (right) stage 9 oocytes. Loss of Ens or Khc induces a strong decrease of Patronin-YFP at the anterior cortex. **(D)** Scatter dot plot showing the mean (± sd) Patronin-YFP intensity at the anterior cortex, in control (248.3 ± 60.55, *n =* 15), *ens* mutant (157.5 ± 60.59, *n =* 13) and *Khc* RNAi (82.96 ± 33.53, *n =* 14) stage 9 oocytes. ***: p<0.00l, ****: p<0.00l (Kruskal-Wallis and Dunn’s tests). **(E)** Shot-GFP cortical localization in a control (left) or in an *ens* mutant (right) stage 9 oocyte. **(F)** Scatter dot plot showing the mean (± sd) Shot-GFP cortical intensity at the anterior cortex, in control (54.26 ± 46.37, *n =* 18) or *ens* mutant (23.88 ± 28.12, *n =* 39). *: p<0.05 (Mann-Withney test). **(G)** Shot-YFP cortical localization in a control (left) or a *Khc* RNAi (right) stage 9 oocyte. **(H)** Scatter dot plot showing the mean (± sd) Shot-YFP mean intensity at the anterior cortex in control (55.09 ± 37.12, *n =* 9) and *Khc* RNAi (19.47 ± 32.46, *n =* 10) stage 9 oocytes. *: p<0.05 (Mann-Withney test). **(I)** Schematic representation of ncMTOCs (magenta) distribution in a control (left) and an ens or *Khc* RNAi stage 9 oocyte (right). For all images, scale bars: 10 µm.

Finally, to investigate the Ens/Khc protein complex contribution to ncMTOC formation at the cell cortex oocytes, we quantified the cortical recruitment of Patronin (Figures 3C and 3D) and Shot (Figures 3E-3H) in *ens* and in *Khc* RNAi oocytes. Patronin and Shot signals were significantly reduced in *Khc* RNAi and *ens* conditions compared to control cells (Figures 3C-3I), demonstrating the contribution of both Khc and Ens to ncMTOC targeting at the cell cortex.

### *ens* and *Khc* oocytes MT organizational defects are caused by decreased ncMTOC cortical anchoring

A previous study has shown that the cortical ncMTOC components Shot and Patronin are required for MT cortical organization (Nashchekin et al., 2016). As *ens* and *Khc* RNAi oocytes exhibits lower ncMTOCs, we examined in details MTs length and organization, as well as yolk vesicle distribution at stage 9 when ncMTOCs are recruited and at stage 10B when the advection step mediated by sub cortical MT twisters occurs. During stage 9, the average subcortical MT length, taken at one focal plane in live oocytes (see methods), was around 6 μm in controls and the MTs exhibited an orthogonal orientation relative to the anterior cortex (Figures 4A left panels, 4C and 4E). This average MT length was similar in *Khc* RNAi and *ens* mutant oocyte, however the MTs displayed a parallel distribution relative to anterior cortex (Figures 4A middle and right panels, 4C and 4E). At stage 10B, the average length of the cortical MT, was 13 μm (Figures 4B right panels and 4C). The mobile yolk vesicles exhibited homogenous distribution within the oocyte, indicative of an ooplasm mixing (Figures 4B and 4E). In *Khc* RNAi and *ens* mutant, the average length of cortical MT, was reduced by 50% compared to controls (Figures 4B and 4C). Moreover, these short MTs were positioned at the anterior half of the oocyte while immobile cytoplasmic vesicles were located at the posterior half of the oocyte, generating a stratified phenotype (Figures 4B upper panel and 4E).

**Figure 4:**
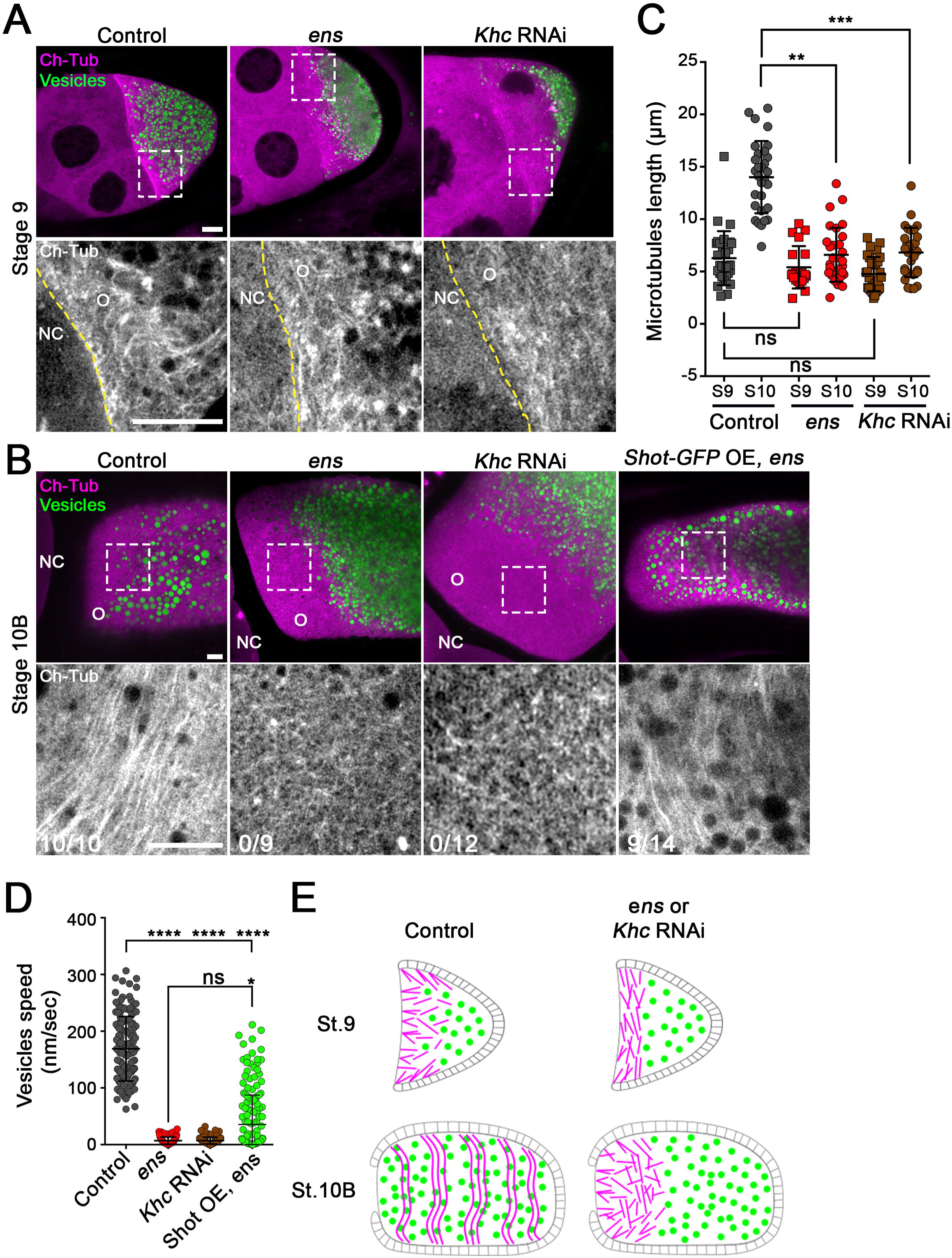
Ens and Khc are required for MT network assembly at stages 9 and 10. **(A)** MT network in control (left), *ens* mutant (middle) and *Khc* RNAi (right) stage 9 oocytes. MTs (Ch-Tub) are shown in magenta and yolk vesicles are shown in green. The white dotted square shows the region shown in the bottom panels. Magnifications (lower panels) show the MT organization (in monochrome). Ens and Khc defective oocytes harbor a parallel MTs distribution while in control MTs are perpendicular to the anterior cortex. **(B)** MT network in control (left), *ens* mutant (middle), *Khc* RNAi (third panel from the left) and in *ens* mutant in which Shot-GFP is overexpressed (Shot-GFP OE, right), at stage 10B. MTs (Ch-Tub) are shown in magenta and yolk vesicles are in green. The white dotted square shows the region shown in the bottom panels. The magnifications views of the cortical MT organization are shown in the lower panels, in monochrome. The number of oocytes presenting long parallel microtubules array to the total number of oocytes observed is indicated at the bottom left. The long cortical MTs that are missing in *ens* mutants (100%, *n =* 10), and are restored following Shot-GFP overexpression in 64% of the oocytes (9/14 oocytes). **(C)** Dot plot showing the mean size (± sd) of cortical MTs in control, *ens* mutant and *Khc* RNAi oocytes during stage 9 and 10B. At stage 9, the mean size of MTs is 5.91 ± 2.79 µm (n = 25) in control oocytes, 4.124 ± 1.20 µm (n = 25) in *ens* oocytes and 4.34 ± 1.43 µm (n = 25) in *Khc* RNAi oocytes. At stage 10B, the mean size of MTs is 14.63 ± 2.05 µm (n = 15) in controls, 6.80 ± 2.43 µm (n = 25) in *ens* oocytes, 5.86 ± 1.72 (n = 20) in *Khc* RNAi oocytes. ns: not significant, **: p<0.0l, ***: p<0.00l. **(D)** Dot plot showing the mean (± sd) vesicle velocities for control (168.9 ± 56.93 nm/sec, *n =* 140) *ens* mutant (6.95 ± 6.45 nm/sec, *n =* 100), *Khc* RNAi (7.01 ± 6.08 nm/sec, *n =* 140) and *ens* mutant overexpressing Shot-GFP (35.48 ± 51.87 nm/sec, *n =* 180) during stage 10B. ns: not significant, *: p<0.05, ****: p<0.00l. **(E)** Schematic representation of the MT network (magenta) and yolk vesicle distribution (green) in control oocytes (left) and *ens* mutant or *Khc* RNAi (right) at stage 9 (top) or stage 10B (bottom). For all images, scale bars: 10 µm. For all plots Kruskal-Wallis and Dunn’s tests were used.

To investigate whether the presence of these short microtubules in *ens* and *Khc* RNAi oocytes is due to defects in microtubule anchoring or is also associated with a decrease in microtubule polymerization at their plus ends, we quantified the velocity and number of EB1 comets, which label growing microtubule plus ends (Figure S3). MT polymerization was not decreased in *ens* and *Khc* RNAi backgrounds (Figure S3), suggesting that the short MTs phenotype detected in *ens* and *Khc* RNAi oocyte was not the consequence of a defective MT polymerization at their plus ends.

Altogether, the analyses of Ens and Khc defective oocytes suggested that MT organization defects during late oogenesis (stage 10B) were caused by an initial failure of ncMTOCs to reach the cell cortex. Therefore, we hypothesized that overexpression of the ncMTOC component Shot could restore certain aspects of microtubule cortical organization in *ens-* deficient oocytes. Indeed, Shot overexpression (Shot OE) was sufficient to rescue the presence of long cortical MTs and ooplasmic advection in most ens oocytes (9/14), resembling the patterns observed in controls (Figures 4B right panel and 4D). Moreover, while *ens* females were fully sterile, overexpression of Shot was sufficient to restore that loss of fertility (Table S1). Altogether, our results strongly support that Ens and Kinesin-1 are required for the cortical recruitment of ncMTOCs, a prerequisite for assembly of the anchored MT streams that sustain ooplasm advection.

Last, to test the possibility that cortical targeting of ncMTOCs by Ens/Khc was possibly MT independent, we treated wild-type oocytes with the MT depolymerizing drug colcemid. We found that these oocytes exhibited lower levels of cortical Shot and Patronin (Figure S4) highlighting the contribution of an intact MT network for Ens and Khc-dependent ncMTOCs localization in the oocyte.

### Interaction of Ens with MTs and Khc are essential for MT organization during oogenesis

To further dissect the contribution of Ens functional domains for the building of the oocyte MT cytoskeleton, Ens^WT^, Ens^LowMT^ and Ens^MutKhc^ GFP-tagged variants were expressed in *ens* mutants and 3 proxies of MT organization and functionality were analyzed in stage 10B oocytes (Figure 5): the presence of long cortical MTs (Figures 4B and 5A), the oocyte polarization by quantifying the recruitment of the Staufen protein to the posterior cortex (Figures 5B and 5C), and the measurement of cytoplasmic advection (Figures 5D and 5E). By contrast to controls, *ens* oocytes lacked long cortical MTs twisters (Figure 4B). These oocytes were also deficient for the recruitment of Staufen-mCherry at the posterior cortex (Figures 5B and 5C) and did not exhibit cytoplasmic advection (Figures 5D and 5E), in agreement with previous studies (Lu et al., 2018; Metivier et al., 2019; Sung et al., 2008). Rescue analyses revealed that Ens^WT^ was able to rescue the presence of long cortical MTs twisters, full Staufen posterior localization and cytoplasmic advection (Figures 5A-5E, (Metivier et al., 2019)). In Ens^LowMT^ oocytes, the long cortical MTs were sparse, Staufen-mCherry posterior recruitment was decreased, and the cytoplasmic advection was lowered compared to Ens^WT^ oocytes (Figures 5A-5E). Finally, Ens^MutKhc^ expressing oocytes, unable to bind Khc, were indistinguishable from *ens* or even from *Khc* RNAi oocytes: they did not show any long cortical MTs twisters, exhibited low amount of Staufen-mCherry protein at the posterior oocyte cortex and no cytoplasmic advection (Figures 5A-5E). Altogether, our result show that Ens binding to MT and to Khc are critical for MT organization.

**Figure 5:**
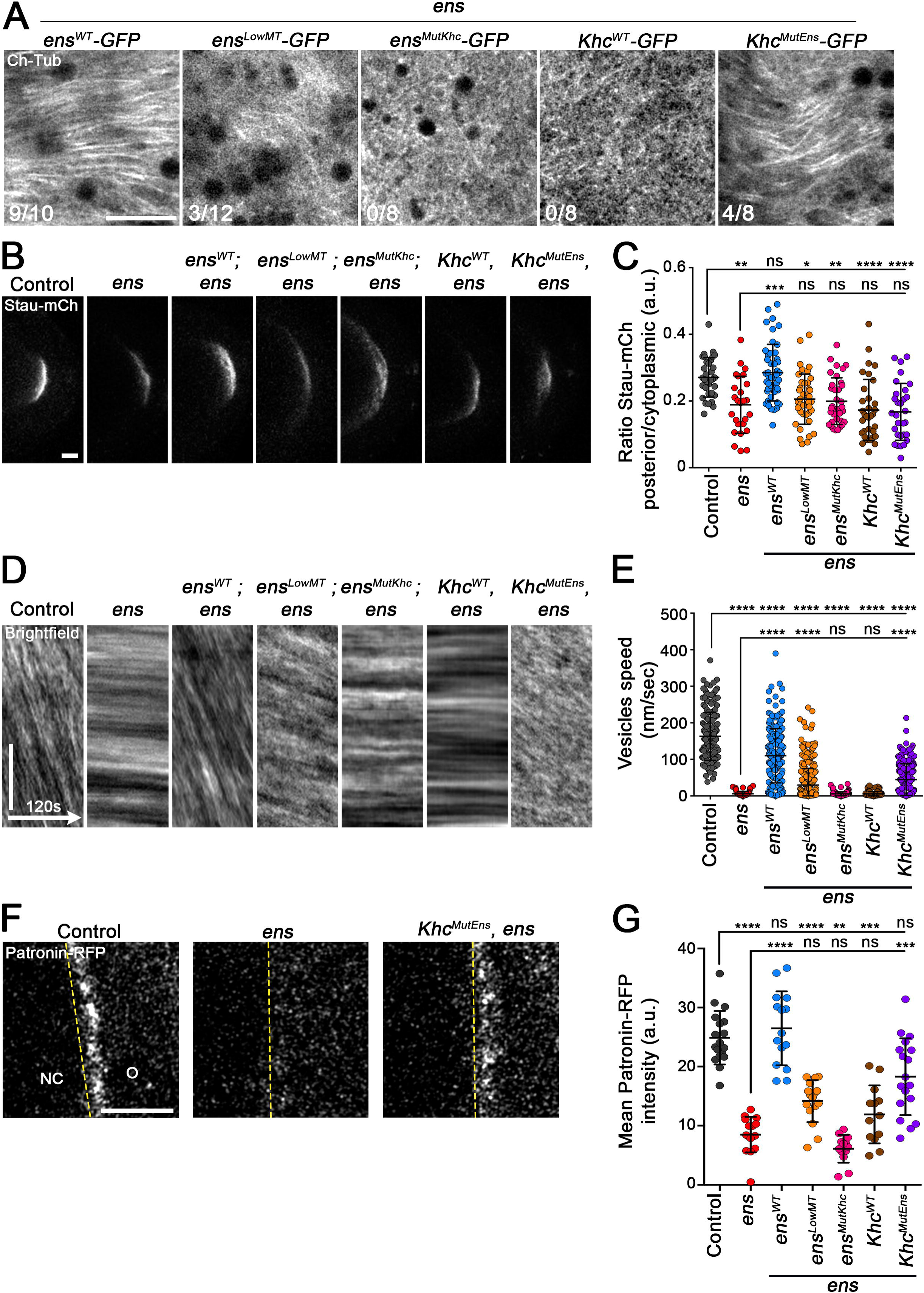
Ens binding promotes relief of Khc inhibition, the formation of the cortical MTs streams that sustains cytoplasmic advection and oocyte polarization. **(A)** Cortical MT cytoskeleton at stage 10B in *ens* oocytes expressing Ens^WT^-GFP, Ens^LowMT^-GFP, Ens^MutKhc^-GFP, Khc^WT^-GFP or Khc^MutEns^-GFP. Ens^WT^-GFP rescues the formation of MTs streams (9/10 oocytes), whereas Ens^MutKhc^-GFP (0/8 oocytes) and Khc^WT^-GFP (0/8 oocytes) did not. MTs streams were also detected in 25% of Ens^LowMT^-GFP (3/12 oocytes) and in 50% Khc^MutEns^-GFP (4/8 oocytes) indicating a partial rescue of the ens phenotype. **(B)** Localization of the Staufen-mCherry crescent at the posterior pole at stage 10B of control, *ens,* and *ens* oocytes expressing Ens^WT^-GFP, Ens^LowMT^-GFP, Ens^MutKhc^-GFP, Khc^WT^-GFP or Khc^MutEns^-GFP. **(C)** Dot plot showing the mean (± sd) posterior/cytoplasmic Stau-mCherry intensity ratio in controls, *ens,* and *ens* mutant expressing each of the indicated GFP tagged protein. Control oocytes (0.27 ± 0.06, *n =* 33), *ens* (0.19 ± 0.09, *n =* 26), *ens* mutant expressing Ens^WT^-GFP (0.29 ± 0.08, *n =* 47), Ens^LowMT^-GFP (0.21 ± 0.07517, *n =* 41), Ens^MutKhc^-GFP (0.20 ± 0.07, *n =* 35), Khc^WT^-GFP (0.17 ± 0.09, *n =* 33), Khc^MutEns^-GFP (0.17 ± 0.09, *n =* 30). **(D)** Kymographs showing displacement of vesicles for 120s in control, *ens,* and *ens* mutant expressing each of the indicated GFP tagged protein. **(E)** Dot plot showing the mean (±sd) vesicle velocities for the indicated genotypes. Control oocytes (163.2 ± 65.85 nm/sec, *n* = 200), *ens* (5.89± 6.36 nm/sec, *n* = 100), *ens* mutant expressing Ens^WT^-GFP (110.0 ± 74.24 nm/sec, *n =* 210), Ens^LowMT^-GFP (29.65 ± 45.16 nm/sec, *n =* 360), Ens^MutKhc^-GFP (5.79 ± 5.50 nm/sec, *n =* 160), Khc^WT^-GFP (5.73 ± 6.05 nm/sec, *n =* 220), Khc^MutEns^-GFP (44.65 ± 43.26 nm/sec, *n* = 240). The expression of Ens^WT^-GFP in *ens* mutant rescued the vesicle movement similarly to controls, whereas Ens^LowMT^ and Khc^MutEns^ partially rescued their speed. **(F)** Localization of Patronin-RFP at stage 9 in control (left), *ens* (middle) and *ens* oocyte expressing Khc^MutEns^-GFP (right panel) conditions. The expression of Khc^MutEns^-GFP restores the localization of Patronin-RFP in an *ens* mutant context. **(G)** Scatter dot plot showing mean Patronin-RFP intensity (±sd) at the anterior cortex in control, *ens* mutant and *ens* mutant expressing Ens^WT^-GFP, Ens^MutKhc^-GFP, Ens^LowMT^-GFP, Khc^WT^-GFP or Khc^MutEns^-GFP stage 9 oocytes. Control oocytes (24.98 ± 4.77, *n* = 17), *ens* mutant (7.63 ± 2.53, *n =* 16), *ens* mutant expressing Ens^WT^ (22.23 ± 8.00, *n =* 11), Ens^LowMT^ (15.64 ± 2.35, *n =* 18), Ens^MutKhc^ (6.09 ± 2.35, *n =* 13), Khc^WT^ (11.91 ± 4.01, *n =* 13) Khc^MutEns^ (19.08 ± 5.73, *n =* 16). Khc^MutEns^ partially rescue the anterior accumulation of Patronin. For all images, scale bars: 10 µm. For all plots, ns, not significant p>0.05; * p<0.05; ** p<0.0l; *** p<0.00l; **** p<0.000l; Kruskal-Wallis and Dunn’s tests were used.

### Relief of Khc auto-inhibition by Ens is crucial for MT organization

The current model of Kinesin-1 activation proposes that Khc exist in an inactive, folded, closed auto-inhibited conformation (Tan et al., 2023a; Weijman et al., 2022; Yildiz, 2024). Binding to accessory proteins, cargoes and regulatory proteins would release this auto-inhibitory state (Hooikaas et al., 2019; Metivier et al., 2019). Based on *in vitro* experiments with truncated proteins, it was hypothesized that a crucial motif in Khc mediates both autoinhibition and physical interaction with MAP7/Ens, subsequently leading to the relief of Khc autoinhibition (Hooikaas et al., 2019; Monroy et al., 2018). So, we hypothesized that the Ens-binding motif in Khc could be relevant for regulating Kinesin-1 activity *in vivo. We* therefore built a full-length Khc^MutEns^ variant in which the Ens-binding motif was mutated (Figure S5A). This variant expressed in *E. coli* displayed reduced Ens-binding properties compared to the WT Khc protein (Figure S5B). However, both Khc^WT^-GFP and Khc^MutEns^-GFP expressed in null *Khc* germline clones were able to restore female fertility indicating both proteins were active in this context (Table S1). When expressed in an *ens* mutant oocyte, the Khc^WT^-GFP transgenic protein, was not able to restore oocytes defects such as the formation of cortical MTs streams, Staufen posterior recruitment or cytoplasmic advection, confirming the importance of Ens for Khc function (Figures 5A-5E). By contrast, expression of the Khc^MutEns^ variant, despite the absence of Ens, was able to restore the formation of some cortical MTs and a slow but significant cytoplasmic advection (Figures 5A-5E). Strikingly, this partial rescue was correlated with a significant increase of the ncMTOC protein Patronin to the cell cortex during stage 9 confirming the strong correlation between ncMTOCs cortical targeting and establishment of MTs twisters later on (Figures 5F and 5G). Similarly, in the context of the fly neural stem cell, Khc^MutEns^-GFP but not Khc^WT^-GFP was able to restore the compromised centrosome separation of *ens* single and in *Khc;ens* double mutants (Figures S5C and S5D). Interestingly, when tested in the same conditions, a previously described active Khc variant with impaired loss of auto-inhibition was also able to rescue centrosome separation (Figures S5D, (Barlan et al., 2013; Friedman and Vale, 1999; Kelliher et al., 2018)). Altogether, these results show that the Khc^MutEns^ variant is active and is able to compensate the absence of Ens in two different cell contexts, highlighting the importance of Ens in relieving Khc auto-inhibition for several crucial Kinesin-1 function during development.

## Discussion

In *Drosophila,* the transition from mid- to late oogenesis marks a pivotal phase where a specialized cortical microtubule network is established. This network plays a crucial role in driving cytoplasmic advection and oocyte polarization. These two processes, essential for ensuring proper embryonic development and precise tissue patterning are regulated by Kinesin-1. In this study, we unveil the molecular mechanisms that govern the spatial activation of Kinesin-1 by its key activator, Ens. We also reveal a critical function of the Ensconsin/Khc complex for the building of its own MT network *via* the cortical recruitment of ncMTOCs.

### MT advection failure in *Khc* and *ens* in late oogenesis stems from defective cortical ncMTOCs recruitment

Ooplasmic streaming, also referred to as cytoplasmic advection or mixing, depends on the precise organization of a cortical microtubule network during late-oogenesis. The mechanisms underlying MT assembly and their role in sustaining advection have long remained unclear. However, recent advances in biophysics, mathematical modeling, and live imaging have provided compelling evidence that advection is driven by cortically anchored MTs. Kinesin-1 recruited to these MTs facilitates the efficient transport of cargoes toward MT plus ends and generating a flux. This cytoplasmic flux at the scale of a single MT induces neighboring MTs bending, leading to an overall collective MT bending that produces robust ooplasmic flows (Dutta et al., 2024; Monteith et al., 2016). In control oocytes, these long, aligned MTs streams are prominent during late oogenesis but are missing in *ens* and *Khc* RNAi oocytes. Instead, our MT network analyses reveal the presence of numerous short MTs cytoplasmic clustered in an anterior pattern. This phenotype prompted us to investigate two potential defective mechanisms: (i) a failure in MT anchoring at the cell cortex possibly during early oogenesis and (ii) an impaired MT growth.

MT anchoring is a process that starts during early oogenesis and is dependent upon ncMTOCs recruitment at the anterior cortex. These ncMTOCs are composed of the actin-binding Spectraplakin homolog Shot and the MT minus-end binding protein Patronin (Nashchekin et al., 2016; Tillery et al., 2018). Our live imaging observations of Shot and Patronin in *ens* and *Khc* RNAi oocytes revealed a marked decrease in their localization at the latero-anterior cell cortex during mid-oogenesis. This low cortical recruitment of ncMTOCs is consistent with poor MT anchoring and their cytoplasmic accumulation, and explain why posterior localization of fate determinants and advection does not occur normally in ens and *Khc* mutant oocytes at later stages. Initially, we hypothesized that the presence of short MTs in late-stage oocytes was the consequence of a defective MT growth. Indeed, previous studies have shown that Ens promotes MT polymer formation both *in vitro* and in mitotic neuroblasts (Gallaud et al., 2014; Thomas et al., 2021). However, MT growth is not significantly impaired in *ens* mutant, suggesting that Ens-mediated MT polymerization is not the key factor in this context. This also corroborates that another MT polymerase, minispindles, the homologue of human chTOG, regulates MT growth in oocytes (Lu et al., 2023). An indirect argument allows us to likely exclude the hypothesis of a role for Ensconsin in MT polymerization: the loss of Khc or Ens produces the same phenotype. As Khc, has not so far been characterized as a protein that stimulates MTs polymerization, the presence of short, unanchored MTs is not directly linked to defective MTs polymerization. Finally, our complementation assays using an Ens variant (Ens^MutKhc^) with intact polymerization activity but defective Khc binding completely fails to rescue the prevalence of short, unanchored cytoplasmic MTs. These findings indicate that Khc activation by Ens, but not Ens ability to promote MTs polymerization, is the critical factor driving both MTs stability and anchoring. Therefore, we propose the following hypothesis: unanchored MTs by ncMTOCs are instable and this phenomenon aligns with previous observations. First, free MT minus ends, obtained following Patronin depletion, depolymerize in multiple cell types, including *Drosophila* S2 cells (Goodwin and Vale, 2011) and dendrites, respectively (Feng et al., 2019). Similarly, reduced cortical MT density has been reported in *Patronin* mutant oocytes, paralleling our observations in *ens* and *Khc* mutants characterized by weak cortical Patronin recruitment, defective MT anchoring and stability. Together, our data suggest that the defective MT-dependent mechanisms observed in *ens* mutant and *Khc* RNAi late oocytes stem from impaired cortical recruitment of ncMTOCs at earlier stage. Supporting this model, an increase of Shot levels, in an *ens* mutant oocyte restored cortical Patronin recruitment, MTs anchoring and growth, enabling cytoplasmic advection. Strikingly, Shot overexpression even rescues female fertility. This strong genetic interaction underscores the tight coupling between MT anchoring and elongation. It also reveals that Ens primary function is to target ncMTOCs to the cell cortex. This rescue suggests that *ens* and *Khc* mutant oocytes are not inherently incapable of hosting ncMTOCs, but rather exhibit defects in their targeting to the cell cortex. An active transport of cytoplasmic ncMTOCs by Khc directly on MTs or indirectly, through cytoplasmic flows likely supports this transport. We believe such MTs and Khc-dependent transport is necessary in this system because oocytes are large cells in which the ooplasm’s high viscosity hinders passive diffusion of large protein complexes over long distances. This model is also supported by three findings. First of all, although a previous study reported that MTs are not required for ncMTOCs targeting to the cortex (Nashchekin et al., 2016), our experiments using a more stringent MTs depolymerization protocol revealed reduced cortical localization of Shot and Patronin, implicating the MT network in ncMTOCs localization. Second, an Ens^MutKhc^ variant, defective in Khc activation, fails to complement Ens-dependent ncMTOCs recruitment to the cortex. Third, a dominant-active Khc^MutEns^ variant partially rescues Patronin localization at the cortex in the absence of Ens. Collectively, these observations strongly support that Kinesin-1-mediated cargo transport is crucial for ncMTOCs cortical recruitment. How ncMTOCs are transported to the cortex is unclear. Similarly to Ens (this paper), Patronin-bound MTs are able to travel from NC toward the oocyte troughs ring canals a place where Shot is present at elevated levels (Lu et al., 2021). It is therefore possible that ncMTOCs/Ens-bound MTs are fully assembled as they cross the ring canals. In the oocyte, these MTs would be fully competent to be transported directly through Khc-dependent MT sliding (Lu et al., 2016) promoting their anchoring to the actin cortex. Alternatively, a local ooplasm flow powered by Khc could also contribute to the ncMTOCs/MTs trapping by the cortical actin cytoskeleton.

### A dual Ensconsin oocyte enrichment mechanism achieves spatial relief of Khc inhibition

Kinesin-1 is an essential motor involved in many physiological processes and many mutations in Kif5A are causal of human diseases (Kawaguchi, 2013). Deciphering the molecular insight of Khc activation is therefore a crucial issue. In the egg chamber, Khc is evenly distributed between the NC cytoplasm and the ooplasm but only active in the oocyte. This have raised two long standing questions: how is Khc activated? Why only in the oocyte? Several answers to these questions are suggested from studies of purified proteins. Recombinant full length Khc is not active without addition of several co-factors in motility assays (Chiba et al., 2022). By contrast, truncated versions of Khc harboring the motor domain but lacking C-terminal regions are actively recruited and move processively on MTs (Monroy et al., 2018). In agreement with these regions exhibiting auto-inhibition properties, structural studies have revealed that Khc inactive heterodimers exhibit a closed/locked conformation associated with multiple intra-molecular interactions between the motor domain and other motifs (Tan et al., 2023b; Weijman et al., 2022). One Khc auto-inhibition motif has been characterized *in vitro:* it is positioned at the C-terminal part of the second coiled-coil dimerization domain. This motif is adjacent to a hinge supposed to confer the flexibility that promotes the closed auto-inhibited state. This motif was also proposed to mediate Ens/MAP7 dependent Kinesin-1 activation and recruitment to MTs (Hooikaas et al., 2019; Monroy et al., 2018). We have mutated this motif in Khc and showed that the resulting Khc^MutEns^ protein loses its interaction with Ens. Strikingly, expression of this Khc^MutEns^ variant was able to partly restore the recruitment of ncMTOCs, the formation of cortical MTs streams, and generate cytoplasmic low speed advection, even in an *ens* null background. In brain neural stem cells, expression of that Khc^MutEns^ restored the defective centrosome separation of *ens* single and *Khc; ens* double mutant neuroblasts, similarly to another active Khc^ΔHinge2^ mutant (Barlan et al., 2013; Friedman and Vale, 1999; Kelliher et al., 2018). Altogether, analyses of Ens/Khc interaction-defective mutants indicates the Ens^MutKhc^ behaves as a null *Khc* allele while the Khc^MutEns^ variant is constitutively active. Our functional analyses demonstrate that Ens binding to Khc therefore promotes relief of Khc auto-inhibition. Despite that Khc^MutEns^ rescued the fertility of *Khc* mutant germline clones’ oocytes, it failed to rescue the development of the fly. As many cargoes transported on MTs are subjected to a tug of war directional motion toward MT plus or minus ends by Kinesin and dynein respectively (Hancock, 2014), this tight equilibrium is probably disrupted in the presence of a constitutively active Khc, detrimental in many cells and for fly development.

How spatial activation of Kinesin-1 is regulated in oocytes? The contribution of aTM1 and Staufen to the transition from Dynein-dependent to Kinesin-dependent cargo transport has recently been described (Gáspár et al., 2023). We now add another piece to the puzzle and address the essential contribution of Ens to the spatial regulation of Khc activity.

The presence of high levels of Ens in the oocyte is making this protein a strong candidate to drive spatial Khc activation in oocytes. When deciphering the molecular mechanisms driving that Ens enrichment in the oocyte, we found that it was MT-dependent; a low MT affinity variant is not enriched in oocytes. Moreover, Ens-bound MTs travel from NC toward the oocyte trough ring canals. Dhc depleted-cells do not enrich Ens in the oocyte. Therefore, by driving transport of Ens-bound MTs in the oocyte, Dynein is an essential regulator of Khc activation. In addition to this, we show that maintenance of high levels of Ens is regulated by Ninein. In muscle and fat body cells, Ninein interacts with Ensconsin, this interaction contributing to modulate the MT network shape in these two cell types. It is also the case in oocytes; *Ninein* mutant oocytes exhibits lower Ens levels. This defect is accompanied by a lower recruitment of Shot at the anterior cell cortex in early oogenesis. This reduced recruitment of ncMTOCs at the cell cortex is nevertheless to generate a cortical MT network and a lower cytoplasmic advection at later stages but in only 50 % of the oocyte. Interestingly, elevating the levels of Ninein promotes the accumulation Ens-containing aggregates at the cell cortex, that also contained Shot and Patronin. Altogether, our results indicate that cortical Ninein levels are intrinsically linked to the recruitment of Ens and ncMTOC components at the cell cortex.

In this study, we have shown that the combination of two mechanisms, a Dynein-dependent transport of Ens-bound MTs, associated with Ens maintenance by Ninein in the oocyte, are needed for the enrichment of Ens in the oocyte. This Ens enrichment activates Khc and promotes the cortical recruitment of ncMTOCs during mid-oogenesis. The data obtained in this study suggest that Kinesin-1 is not only a mere user of the microtubule network but also a key regulator of its structural foundations.

## Acknowledgments

We thank, Christelle Benaud, Gregory Eot-Houllier, Anne Pacquelet, Chantal Roubinet, Laurent Richard-Parpaillon for critical readings of the manuscript. We thank Antoine Guichet, Daniel St Johnston, Xin Liang, Mary Baylies, Timothy Megraw, David Glover, Anne Ephrussi, Jill Wildonger, Renata Basto, Kassandra Ori-McKenney, Eric Lecuyer, Romain Gibeaux and Emmanuel Derivery for their kind gift of reagents. We thank Helene Bouvrais for help in computer image analysis. We thank the Bloomington and Kyoto Stock Centers for *Drosophila* strains and the Drosophila Genome Research Center for plasmids. We acknowledge Stéphanie Dutertre, Gilles Le Marchand and Xavier Pinson of the Microscopy Rennes Imaging Center (MRic, BIOSIT, Biogenouest) for assistance in microscopy.

## Materials and Methods

### Fly stocks and genetics

All *Drosophila melanogaster* stocks were grown on standard medium at 25°C. Strains used were: *FRT42B, Khc^27^* (obtained from Guichet A., Institut Jacques Monod, UMR 7592, Paris, France, (Januschke et al., 2002)), *Ubi-Ens-GFP* and *Ubi-Eb1-EGFP* (Gallaud et al., 2014), *Patronin-YFP, shot-YFP, UASp-Cherry-Patronin* (gift from St Johnston D., The Gurdon Institute and the Department of Genetics, Cambridge, UK, (Nashchekin et al., 2021; Nashchekin et al., 2016)), *Patronin^RFP-KI^* (gift from Liang X., Tsinghua-Peking Joint Center for Life Science, School of Life Sciences, Tsinghua University, Beijing, China, (Sun et al., 2021)), *Ninein^null^ (Bsg25D^null^;* gift from Baylies M., Sloan Kettering Institute, New York, USA, (Rosen et al., 2019)), *UAS-Ninein-TagRFP-Myc* (gift from Megraw T.L., Department of Biomedical Sciences, College of Medicine, Florida State University, Tallahassee, USA, (Tillery et al., 2024)), *Ubi-pavarotti-EGFP* (gift from Glover D., Department of Genetics, University of Cambridge, UK, (Minestrini et al., 2002)), *Khc^63^* (gift from Ephrussi A., (European Molecular Biology Laboratory, Heidelberg, Germany, (Djagaeva et al., 2012)), *Khc^ΔHinge2^* (gift from Wildonger J., Biochemistry Department, University of Wisconsin-Madison, Madison, WI, USA, (Kelliher et al., 2018)), *Ubi-RFP-αTub* (gift of Basto, R., Institut Curie Paris, France, (Basto et al., 2008)).

Experiments were performed in trans heterozygous *ens^Δ3277^ /ens^ΔN^* flies previously described (Metivier et al., 2019; Sung et al., 2008) and referred as *ens* mutant. The following strains were obtained from the Bloomington Drosophila Stock Center: *ens^Δ3277^* (BDSC: 51319), *ens^ΔN^* (BDSC: 51317). *matα4-GAL-VP16-V37* (BDSC: 7063), *matα4-GAL-VP16-V2H* (BDSC: 7062), *UAS-mCherry-Tub* (BDSC: 25774), *insc-GAL4; UAS-mCherry-Tub* (BDSC: 25773), *Ubi-GFP* (BDSC: 1681), *UAS-shot-GFP* (BDSC: 29042). *matα4-V2H* was used to drive all UAS constructs except for *UAS-Ch-Tub* (Figure S2B) drived by *matα4-V37*.

The RNAi lines used were: TRiP.GL00543 (BDSC: 36583) for *Dhc64c* and TRIP.GL00330 (BDSC: 35409) for *Khc.* Expressions of RNAi with *matα4-GAL-VP16-V2H* were carried out at 29°C from pupa to adult stages.

Germline clones of *FRT42B, Khc^27^* were generated by the FLP/FRT system (Chou et al., 1993; Chou and Perrimon, 1996), by incubating L2-L3 larvae in a 37°C water bath for 1 hour on 3 consecutive days, or pupae for two hours during two days. *FRTG13 OvoD1-18* (BDSC: 4434) was used, and homozygous clones were selected using the *OvoD1* system.

### Generation of Khc and Ens variants and expression vectors

Khc open reading frame (clone DGRC SD02406) was PCR amplified and cloned in a gateway pENTR vector without the stop codon to generate a Khc entry clone. The conserved polarized amino acids starting at position E512 (EELAVNYDQK) supposed to mediate interaction with HsMAP7 (Hooikaas et al., 2019; Monroy et al., 2018) were mutated into AALAVNYAAA to generate a Khc^MutEns^ variant defective for Ens binding (Figures S5A and B).

The Ens entry clone was described before (Gallaud et al., 2014). The conserved amino-acid stretch in Ens from position 690 (REERRKR) required for Khc activation (Monroy et al., 2018) were mutated into 7A to produce an Ens^MutKhc^ variant with defective Khc-binding properties (Figure S1C, bottom). Human MAP7/HsEnsconsin binding to MT is downregulated following Cdk1 phosphorylation on S and T located on the N-terminal MT-binding domain (McHedlishvili et al., 2018). This *in vivo* modulation of MT binding was used to generate an Ens variant with lower MT affinity (Ens^LowMT^) by mutating (T144, S158, S213, T282, S288, T316) into D (Figure S1C, middle). The Ens^LowMT^ entry clones were obtained using the QuickChange lightning multi-site-directed mutagenesis kit (Thermo Fisher Scientific).

The pUWG-Ens and pET102-Ens vectors were described before (Gallaud et al., 2014) and were used as backbone vectors to produce Ens^MutKhc^ and Ens^LowMT^ for fly and bacterial expression respectively. The Khc ORF and poly Ubiquitin promoter were cloned into pUAS-K10-attB-GFP vector to generate the fly pUbi-attB-Khc-GFP expression vector. Khc was cloned into pET28-mScarlett-Strept (Monroy et al., 2018), a kind gift of K. Ori-McKenney to produce an *E.coli* expression vector. pUbi-attB-Khc-GFP and pET28-Khc-mScarlett-Strept were used as backbone vectors to produce the Khc^MutEns^ expression vectors. Ens and α-tubulin 84B were tagged with Dendra2 and InfraRed Protein respectively into pUAS-K10-attB fly expression vector (Kachaner et al., 2017). Unless specifically mentioned, all cloning procedures were performed using the In-Fusion cloning kit (Takara).

### Generation of transgenic flies

pUWG vectors were injected into *w^1118^.* pUAS-K10-attB-IRP-α-Tubulin vectors were injected into flies using the VK02 (BDSC: 9723) and VK27 (BDSC: 9744) landing sites. UAS-K10-attB-Dendra2-Ens were injected into VK27 (BDSC: 9744). pUbi-attB-Khc^WT^-GFP and pUbi-attB-Khc^LowEns^-GFP were injected into the VK05 landing site (BDSC: 9725) by BestGene Inc.

Endogenous mCherry tagging of the *Staufen* gene was performed by CRISPR/Cas9. Staufen::mCherry lines were homozygous viable. All injections of plasmids were performed by BestGene Inc.

### Expression and purification of recombinant protein

The pET102, pDest42, pET28 and pET23b expression vectors were transformed into the *E. coli* BL21(DE3) strain (New England Biolabs). The recombinant protein expression was induced with 1mM IPTG during 4 h at 25°C for pET102 vectors and overnight at 16 °C for pET82 vectors. The bacterial pellets were stored at -20°C until use. The pellets were resuspended on ice, in Lysis Buffer (LB: 200 mM Tris-HCl (pH 7,5), NaCl 1 M, 10% glycerol, 10 mM Imidazole, lysozyme x1000 for pET102, pET23b and pET23b vectors or LB: 50 mM NaH_2P_O_4_, 300 mM NaCl, 10% glycerol, lysozyme x1000, 1 mM DTT for pET28 vectors containing 1% Triton X-100 (Sigma), proteases inhibitor tablets (Roche) and lysozyme (1mg/ml). The resuspended bacterial pellets were then sonicated 10 times for 10 s on ice and centrifuged at 10 000 *g* for 30 min at 4°C. The 6xHistidine-tagged WT and mutant Ens proteins and Streptavidin-tagged WT and mutant Khc variants were purified following the manufacturer’s s instructions on Nickel or Strept-Tactin columns (Qiagen). The proteins were then dialyzed overnight in PBS at 4°C, concentrated on Amicon Centrifugal Unit (Millipore).

### MT self-assembly and turbidity assays

A 40 µM tubulin solution, purified from porcine brain, was prepared in 1 mM GTP in BRB80 and centrifuged at 30,000 g for 10 minutes at 4°C prior to polymerization (Weis et al., 2010). Tubulin polymerization was then initiated by incubating the samples at 35°C, with the process being monitored turbidimetrically at 350 nm in quartz cuvettes using a UVIKON XS spectrophotometer (BioTek Instruments). The activities of Ensconsin variants were assessed by adding to tubulin different Ensconsin variants at a fixed concentration of 0,375 µM. To evaluate protein aggregation post-assembly, the temperature was reduced to 4°C after 40 minutes of recording (Weis et al., 2010). The spectrophotometric data were adjusted to account exclusively for microtubule polymerization. A linear regression analysis was applied to the portion of the curve corresponding to the polymerization phase, from minute 1 to 15.

### Immunoprecipitation experiments using purified proteins

Purified recombinant GFP proteins (approximately 15 µg) were incubated with 10 µl of GFP-Trap^®^MA (Chromotek) magnetic beads for 1 h at 4°C with rotation. After a wash covered-beads were incubated with a second purified recombinant protein (approximately 1 µg) for 30 min at 4°C with rotation. The beads were washed three times and the bound proteins were analyzed by Western blotting.

### Immunoprecipitation from ovarian extracts

Ovarian extracts were prepared from GFP-tagged transgenic flies by dissecting ovaries (30 flies per genotype). The ovaries were lysed in 200 µl of lysis buffer [10 mM Tris, 150 mM NaCl, 0.5 mM EDTA (pH 7.4)] supplemented with 0.05% NP-40, 1 mM DTT containing 5× proteases inhibitors (Roche). The extracts were centrifuged at 10,000 g for 30 min at 4°C and the supernatants were incubated with 10 µl of GFP-Trap^®^MA (Chromotek) magnetic beads for 1h at 4°C with rotation. The beads were washed three times and the bound proteins were analyzed by western blotting.

### Western blotting analyses

After transfer of the proteins on a nitrocellulose membrane (GE Healthcare) using a Trans Blot Turbo system (Biorad), the membranes were blocked for 2 h with TBST (20 mM Tris, 150 mM NaCl and 0.1 % Tween20, pH 7.4) containing 10 % skimmed milk. Primary antibody incubation was performed overnight at 4°C in TBST containing 5 % skimmed milk. Incubation with the secondary antibodies was performed in 5 % skimmed milk for 2 h at room temperature. Three 15 min washes in TBST were performed after antibody incubation. For western blotting, ECL reagent (ref. 34095 or 34077) was purchased from Thermo Fisher Scientific.

### Immunofluorescence in fly ovaries

Ovaries were collected from two-day-old females and fixed in 4% Paraformaldehyde, 0,1% Triton for 20 min at room temperature. After washes, ovaries were blocked 1 h in PBS, 0,1% Triton, 5% BSA, and incubated overnight with primary antibodies at 4°C. After washes, ovaries were incubated 1h with the secondary antibodies tagged with Alexa Fluor (1:500) and Phalloidin 550 (1:200, Sigma 19083). Ovaries were mounted on a slide with ProlonGold medium with DAPI (Invitrogen).

Alternatively, to visualize and preserve the MT network (Figure 2 and Figure S4), we adapted a protocol used for fly spermatocytes (Cenci et al., 1994; Gatti et al., 1994). The ovaries were briefly washed in PBS and gently flattened in a drop (15 μl) of PBS containing 5% glycerol between an ethanol washed slide and a 18x18 mm coverslip. The preparation was immediately frozen in liquid nitrogen. The slide was removed from the liquid nitrogen tank using a pair of forceps and the coverslip was flipped off using a scalpel blade. The slides with the frozen ovarioles were immediately fixed for 30 min in -20°c methanol and rehydrated for 30 min PBS. After 3 10 min washes in PBS, the samples were permeabilized with PBS containing 0.5% Triton X-100 for 30 min without agitation and washed for 10 min 3 times in PBS. Primary antibodies were diluted in PBS containing 0.1% Triton X-100 and 1% BSA and the samples were incubated overnight at 4°C. After 3 times 5 min washes in PBS, the samples were incubated with an Alexa-conjugated fluorescent secondary antibodies for 4 hours at room temperature. The ovaries were then washed 3 times for 5 min each time in PBS and mounted in ProLong Gold containing DAPI (Invitrogen). Image acquisition was performed using a Leica SP5 confocal microscope. All image analyses were performed using ImageJ (Rueden et al., 2017). For all images, a single section is presented, except when mentioned.

### Colcemid treatment

To depolymerize MTs, ovaries were dissected in Schneider medium and incubated in Schneider medium containing DMSO or 100 μM Colcemid (Sigma C3915) for 60 min before being processed for fixation and immunofluorescence analyses.

### Antibodies

Polyclonal rabbit anti-Ens antibody was used 1:2000 for Western blotting (Gallaud et al., 2014). The YL1/2 rat monoclonal anti-tyrosinated tubulin antibody (ref. Mab1864, 1:200). The anti-GFP mouse monoclonal antibody mix (ref. 11814460001, 1:1000) was obtained from Roche. The anti-GFP chicken antibody was from Abcam (ref. AB183734, 1:2000). The mouse monoclonal anti-Shot (ref. AB_528467, 1:10), were purchased from the Developmental Studies Hybridoma Bank (Iowa University). The guinea pig polyclonal anti-Bsg25D/DmNinein (serum B) was used at 1:1000 for immunostaining and at 1:20000 for Western blotting and was kindly given by Eric Lecuyer (University of Montreal (Kowanda et al., 2016)). The rabbit polyclonal anti-Patronin antibody was used at 1:2000 and was kindly provided by Emanuel Derivery (Derivery et al., 2015). The rabbit polyclonal anti-KHC antibody (1:2000 for Wester Blotting and 1:500 for Immunofluorescence) was obtained from Cytoskeleton. Goat secondary peroxidase-conjugated antibodies (1:5000) were obtained from Jackson ImmunoResearch Laboratories, and donkey Alexa Fluor-conjugated secondary antibodies (1:1000) were obtained from Life Technologies.

### Live imaging of ovaries

Fixation methods affect the shape of the MT network and notably the parallel arrays of long MTs streams of stage10B oocytes (Serbus et al., 2005). Therefore, we mainly used live imaging of ovaries carrying fluorescent reporter genes. Ovaries were dissected on a slide in mineral oil (Sigma M8410) and immediately imaged on confocal microscope at 25°C. We used a *matα4-GAL-VP16 (V37* or *V2H), UAS-ChRFP-Tub* or *UAS-InfraRed-Tub* to visualize the MT network in association with GFP or RFP-tagged reporter genes. Auto-fluorescent yolk vesicles were visualized with 405 illumination.

Living ovaries were imaged using a Zeiss LSM 980 Airyscan2 confocal microscope at 40× or 63× objective plus optical zoom, a Leica SP5 or a Leica SPE microscope. For brightfield images used to quantify vesicles velocity, a DMRXA2 microscope (Leica) was used. For all images, a single section is presented, except when mentioned.

### Live imaging of Dendra-Ens photoconvertion

Photoconversion experiments were performed on Zeiss LSM 980 Airyscan2 equipped with a Plan APO 63X/1.4 N.A. oil-immersion objective. Dendra-Ens was photo-converted in NC in front of a ring canal that connects the nurse cell to the oocyte on rectangle of 10 µm width and 5 µm height. Rings canals were located with the Pavarotti-GFP signal (Lu et al., 2021; Minestrini et al., 2002). Images were acquired every 2 seconds during 3 minutes.

### Quantification of GFP fluorescence ratio between the oocyte and NCs

To measure the GFP-tagged protein enrichment in the oocyte, the mean fluorescence ratio between the whole oocyte and the nurse cells cytoplasm was calculated at stage 8.

### Quantification of the GFP-tagged variant localization on MTs

The quantifications of of Ens^WT^-GFP, Ens^LowMT^-GFP and Ens^MutKhc^-GFP on MTs were performed on stage 9 oocytes expressing each GFP construct in a matα4-GAL, UAS-ChRFP-Tub; ens background. Mean intensity of GFP (GFP^MT^) and Cherry (ChRFP^MT^) signals were measured along individual MT, using the Image J segmented line tool with a width of 10. The mean intensity background of GFP (GFP^background^) and Cherry (ChRFP^background^) signals were measured outside of the oocyte, and subtracted from the GFP and Cherry signals. The resulting mean intensity ratios (IR) were calculated. IR = ((GFP^MT^-GFP^background^)/GFP^background^)/((ChRFP^MT^-ChRFP^background^)/ ChRFP^background^).

### Measurements of the minimal cortical MT length in live oocytes

The cortical MT network was imaged on oocytes expressing *matα4-GAL, UAS-ChRFPTub* using a Zeiss Airyscan 2 confocal microscope. MT networks at stages 9 and 10B are dynamic, making multi-plane analysis and 3D MT network reconstruction challenging. So, the MT lengths were measured in a single subcortical plane, with the limitation that some MTs may extend beyond the analysed plane, potentially leading to an underestimation of their actual length. The MTs lengths were manually measured using the segmented line selection tool on Fiji software, on stage 9 and 10B control, *ens,* or *Khc* RNAi oocytes. 5 MTs per oocyte were measured.

### Quantification of Eb1-GFP comet velocity and number

Images of stage 9 oocytes expressing *Ubi-Eb1-GFP,* were acquired every 0,58s for 1min. The background noise was removed after application of a median filter (radius = 15 stack). Kymographs were generated using the Multi Kymograph plug-in on Fiji. Ten kymographs were generated for each oocyte (n> 7 oocytes) to quantified the Eb1-GFP comets velocities.

For quantification of Eb1-GFP comet number, a threshold was applied to isolate signal from comets in the oocyte. Comets were counted using the analyze particle tool on Fiji. The result is expressed in number of comets per 100 µm^2^.

### Quantification of Patronin and Shot signals

The mean fluorescence of Patronin-YFP or Shot-GFP (Figure 4) and Patronin-RFP (Figure 5) were measured on living stage 9 oocytes along the anterior oocyte cortex. The mean fluorescence of Patronin-YFP, Shot and Tub (Figure S4) were measured on stage 9 immunostained oocytes were measured along the antero-lateral cortex. The segmented line selections tool with a width of 20 (Fiji software) were used. The signal was subtracted by the background noise measured outside the oocyte.

### Staufen-mCherry crescent quantification at the posterior cortex

Stage 10 oocytes expressing Staufen-mCherry were imaged every 1 µm. A sum Z-projection of the oocyte was created using Fiji. The polygon selection tool in Fiji was used to measure the total fluorescence intensity at the posterior crescent and in the whole oocyte to calculate the Staufen-mCherry recruitment ratios.

### Visualization and speed of yolk vesicles

Oocytes were imaged with bright-field on a DMRXA2 microscope (Leica). Images were acquired every 2 s for 2 min. Kymographs were generated using the Fiji Multi Kymograph tool. Four kymographs were generated on either side of each oocyte (n = 5 oocyte min), where flows are seen in fast-streaming stage 10B oocytes. The velocities of twenty particles were quantified per oocyte and averaged to determine the streaming rate (Metivier et al., 2019).

### Quantification of dorsal appendage phenotypes

20 to 30 flies, of both sexes in equal proportions, were transferred into egg-laying cages with grape juice agar plates supplemented with fresh yeast paste and incubated at 25° overnight. Embryos laid on the plate are then detached with water and a fine brush, and transferred to a grid plate for manual counting. Embryos are classified into three categories: embryos with two well-separated dorsal appendages, embryos with two glued dorsal appendages or a single dorsal appendage, and embryos without dorsal appendages.

### Analysis of centrosome separation in larval brain neuroblasts

The analysis of centrosome separation in larval brain neuroblasts was performed as previously described (Berisha et al., 2024). Briefly, the brains of L3 larva expressing *Ubi-RFP-αTub* were dissected and mounted in Schneider’s media. Movies of dividing neuroblasts were acquired with a spinning disk system mounted on an inverted microscope (Eclipse Ti; Nikon). Z series were acquired every 30s. In dividing neuroblasts, the two centrosomes were visualized with their microtubule nucleation potentials shortly before mitosis. The separation angle between the two centrosomes was determined 3min before the nuclear envelope breakdown (NEBD), by drawing a line crossing the two centrosomes (A and B) and the nucleus center. The coordinates in space (x, y, z) are collected to calculate the α angle between the A centrosome, the nucleus center, and the B centrosome, as a read out of centrosome separation.

### Statistical analysis

For all experiments, data plots and statistical analyses were performed with GraphPad Prism 8.4. Graphs show all individual values (dots), the mean and the standard deviation (black lines). Mean values and standard deviation (sd) are indicated in figure legends. The statistical tests used are indicated in each figure legend. For statistical comparisons of two datasets unpaired t-test were performed when data passed the Shapiro and D’Agostino & Pearson normality tests and when sample variance was equal, Mann-Whitney test were performed when data did not pass the Shapiro and D’Agostino & Pearson normality tests. For statistical comparisons of more than two datasets Kruskal-Wallis non-parametric test and Dunn’s multiple comparisons were used when data when data did not pass the Shapiro and D’Agostino & Pearson normality tests and the sample variance test. Asterisks show the significance of variation (ns, not significant p>0,05; *: p<0.05; **: p<0.0l; ***: p<0.00l; ****: p<0.000l).

## Figure legend

**Figure S1:**
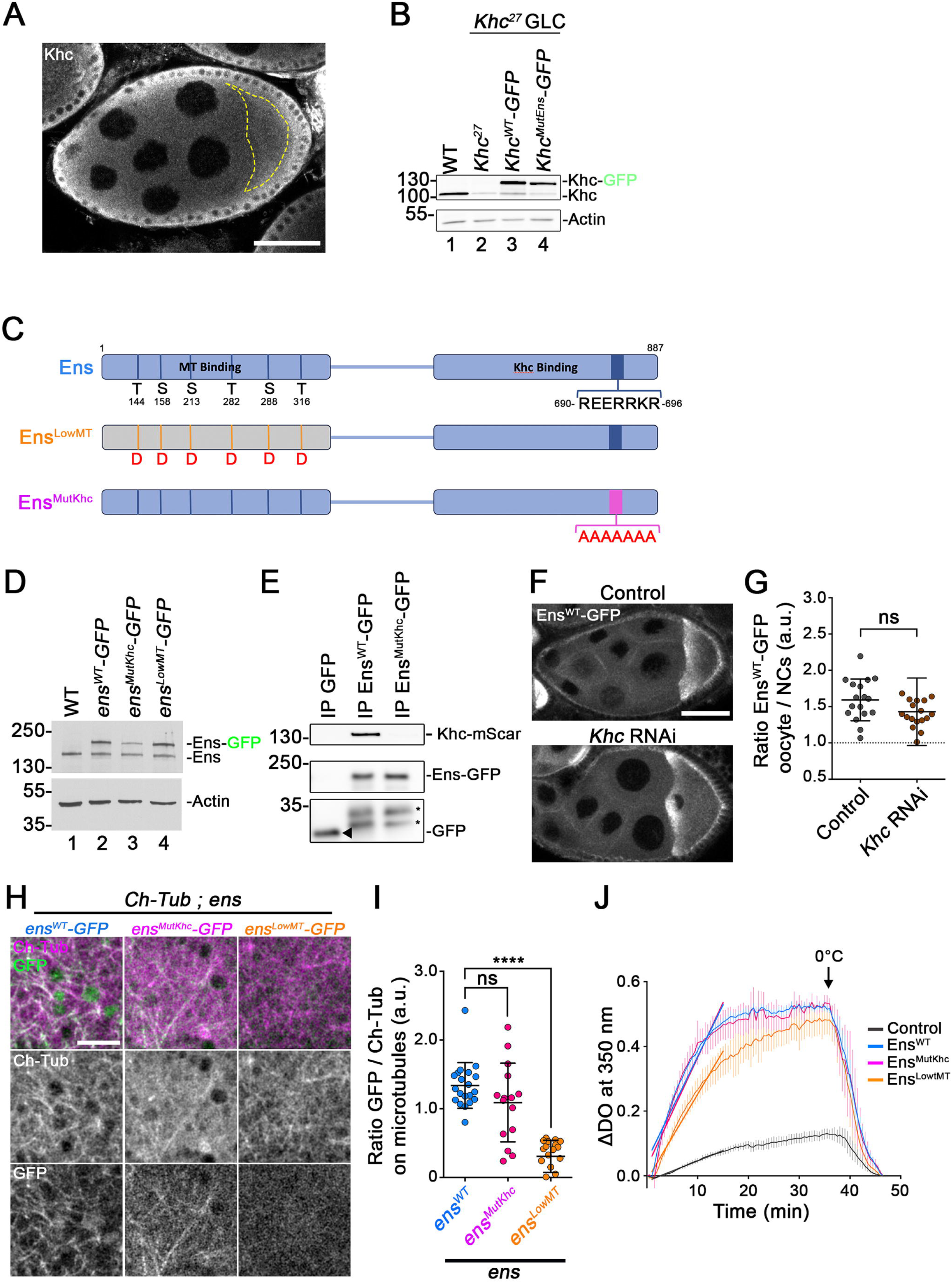
Characterization of Ens mutants. **(A)** Analysis of endogenous Khc localization in a stage 8 egg chamber by immunostaining. Image is 3 µm projection. Scale bar: 50 µm. **(B)** Western blot analysis of Khc, Khc-GFP, Khc^MutEns^-GFP protein levels during oogenesis. Ovaries extracts were probed using anti-Khc (top) and anti-Actin antibodies (bottom) to show the expression levels of endogenous Khc and GFP-tagged Khc constructs in WT extracts (lane 1), *Khc^27^* germ line clones (GLC; lane 2) and *Khc^27^* GLC expressing Khc^WT^-GFP (lane 3) or Khc^MutEns^-GFP (lane 4). **(C)** Schematic representations of Ens and the variants used in this study. Ens harbors a N-terminal MT Binding domain (MBD) and a C-terminal Khc binding domain (KBD). The Cdk1 consensus phosphorylation sites (S and T) and their positions in the MBD are indicated at the bottom (top). The Ens^LowMT^ variant exhibits Cdk1 phospho-mimicking amino-acids substitutions (into D) to lower Ens affinity for MTs (middle). The C-terminal Khc binding region harbors a conserved REERRKR motif (dark blue). That motif is mutated into AAAAAAA (magenta) to generate an Ens^MutKhc^ variant with reduced Khc interaction. **(D)** Western blot analysis of Ens, Ens^WT^-GFP, Ens^LowMT^-GFP, Ens^MutKhc^-GFP during oogenesis. Ovaries extracts were probed using anti-Ens antibodies (top) and anti-Actin antibodies (bottom) and show the expression levels of endogenous Ens and GFP-tagged Ens constructs in WT extracts (lane 1), Ens^WT^-GFP (lane 2), Ens^MutKhc^-GFP (lane 3) and Ens^LowMT^-GFP (lane 4) conditions. The tagged proteins were expressed in a WT background. **(E)** Purified recombinant GFP, Ens^WT^-GFP and Ens^MutKhc^-GFP proteins were immobilized on GFP-TRAP beads and tested for Khc-mScarlet binding. The beads were then analyzed by Western blotting for the presence of Khc (top) or GFP (bottom) with the corresponding antibodies. Khc-mScarlet is present in Ens^WT^-GFP but not in GFP or Ens^MutKhc^-GFP immunoprecipitates. GFP is indicated with an arrow head, * shows degradation fragments of Ens^WT^-GFP and Ens^MutKhc^-GFP. **(F)** Ens^WT^-GFP distribution in a control (top) or a *Khc* RNAi egg chamber (bottom). Scale bar: 50 µm. **(G)** Scatter dot plot showing the mean Ens^WT^-GFP intensity ratio (± sd) between the oocyte and NC in control (1.59 ± 0.29, *n =* 17) and *Khc* RNAi egg chambers (1.43 ± 0.47, *n =* 19). ns, not significant p>0.05 (T-test). **(H)** Localization of Ens^WT^-GFP (left), Ens^MutKhc^-GFP (middle) and Ens^LowMT^-GFP (right) on MTs in stage 9 *ens* mutant oocytes expressing Ch-Tubulin. GFP-tagged proteins and MTs are displayed in green and magenta in the merge panels respectively. Middle and bottom panels show the MTs and GFP-tagged Ens variants in monochrome. Scale bar: 10 µm. **(I)** Scatter dot plot showing the mean GFP to Ch-Tub intensity ratio on MTs (± sd) in stage 9 *ens* oocytes expressing Ens^WT^-GFP (1.34 ± 0.33, *n =* 20), Ens^MutKhc^-GFP (1.09 ± 0.57, *n =* 15), or Ens^LowMT^-GFP (0.31 ± 0.23, *n = 20).* ns, not significant p>0.05; **** p<0.000l (Kruskal-Wallis and Dunn’s tests). **(J)** Analysis of tubulin polymerization *in vitro* by spectrophotometry at 350 nm in the presence of recombinant Ens^WT^, Ens^MutKhc^ or Ens^LowMT^. Tubulin concentration 40 μM, Ens proteins 0,375 μM. Corresponding linear regressions for the phase of polymerization (x = 1 to x = 15) are shown with straight bold lines in controls (n = 4, y = 0,0063x -0,017, r^2^ = 0,9922), Ens^WT^ *(n =* 6, y = 0,0321x + 0,0476, r^2^ = 0,9455), Ens^MutKhc^ (n = 3, y = 0,0339x + 0,0274, r^2^ = 0,9144) and Ens^LowMT^ (n = 4, y = 0,0266x-0,0137, r^2^ =0,9144). Ens^WT^ and Ens^MutKhc^ increased the rate of MTs polymerization and the total number of polymers (blue and magenta curves respectively), compared to the buffer alone (black curve). The Ens^LowMT^ displayed a reduced MTs polymerization rate (orange curve). The slopes of the curves are significantly different; *p < 0.0001* (Linear regression test).

**Figure S2:**
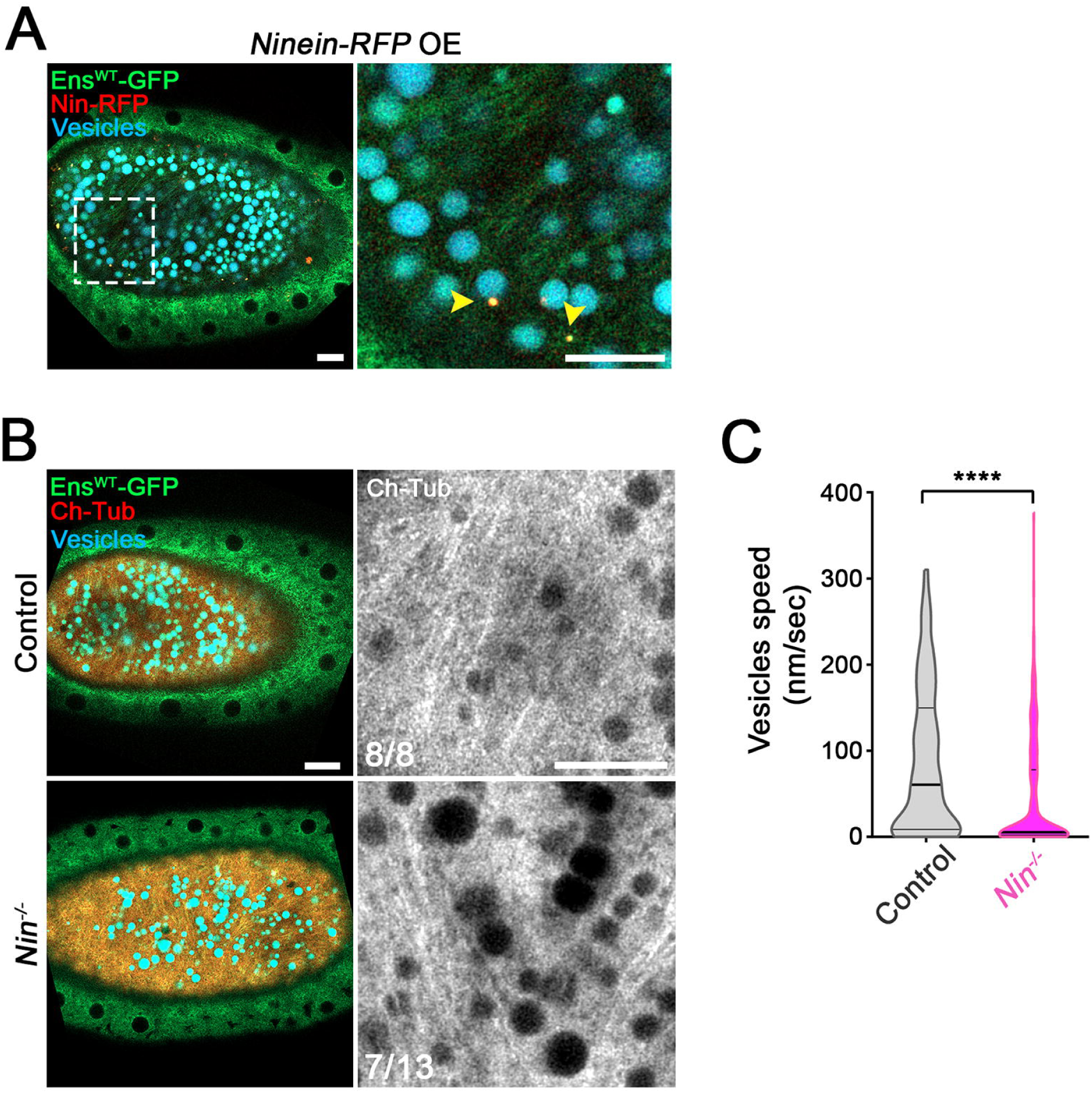
MT network at stage 10B, in Ninein overexpressing and Ninein null oocytes. **(A)** Ninein-RFP (red) and Ens^WT^-GFP (green) cortical aggregates are barely detected during the advection process at stage 10B (right). Magnification (left) of the corresponding region marked by the dotted lines in the right panel, shows few small Ens^WT^-GFP (green) and Nin-RFP (red) cortical aggregates (yellow arrowhead). Ens^WT^-GFP decorates the MT network at this stage. The yolk vesicles autofluorescence is shown in cyan. **(B)** The presence of cortical MTs streams is compromised in *Ninein* mutants. MT network in control (top) and *Ninein* mutant (bottom). MTs (Ch-Tub) are shown in red, yolk vesicles are shown in cyan and Ens^WT^-GFP in green. The magnifications views of the cortical MT organization are shown in the right panels, in monochrome. The number of oocytes presenting long parallel microtubules array to the total number of oocytes observed is indicated at the bottom left. Long cortical MTs are detectable in 53% of *Nin* mutant oocytes. **(C)** Violin plot showing the mean (± sd) vesicle velocities for control (87.12 ± 85.22 nm/sec, *n* = 120) and *Ninein* mutant (62.28 ± 94.54 nm/sec, *n =* 280). **** p<0.000l (Mann-Witney test). For all images, scale bars: 10 µm.

**Figure S3:**
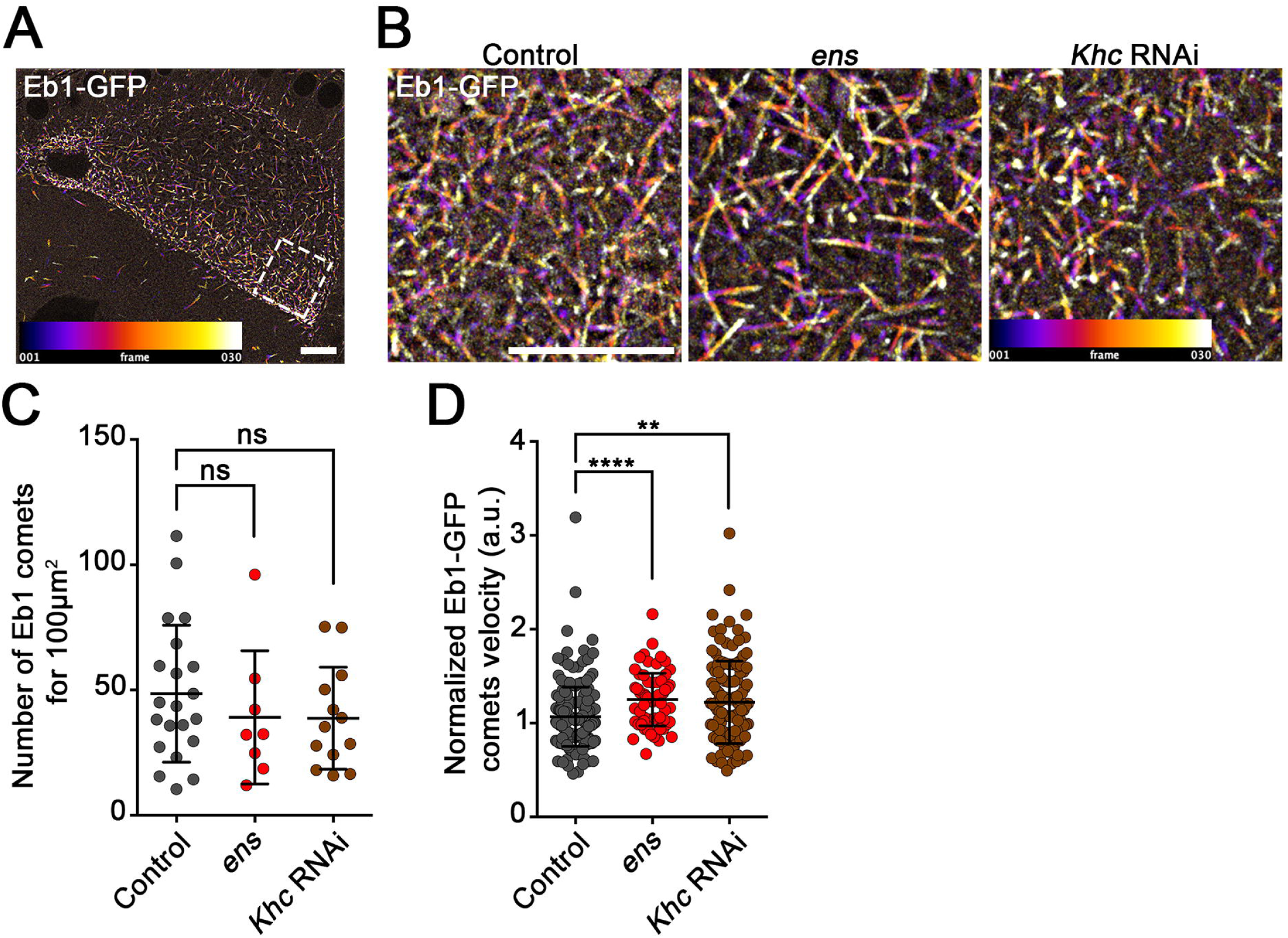
*ens* and *Khc* oocytes do not show decreased MT polymerization at their plus ends. **(A)** Eb1-GFP comet trajectories in a control stage 9 oocyte. Color code indicates the time projection for 30 frames (0,58 s between frames). Region marked by the dotted lines corresponds to the magnification in panel (B). **(B)** Eb1-GFP comet trajectories in control, *ens* mutant and *Khc* RNAi stage 9 oocytes. Color code indicates the time projection for 30 frames (0,58 s between frames). **(C)** Scatter dot plot showing the number (± sd) of Eb1-GFP comets for 100 µm^2^ in control, *ens* mutant and *Khc* RNAi oocytes at stage 9. Control (48.54 ± 27.33, *n =* 21), *ens* (39.11 ± 26.60, *n* = 8), *Khc* RNAi (38.73 ± 20.36, *n =* 13); ns, not significant p>0.05 (Kruskal-Wallis and Dunn’s tests). **(D)** Scatter dot plot showing the Eb1-GFP comets velocity (± sd) in stage 9 control, *ens* mutant and *Khc* RNAi oocytes. Control (1.07 ± 0.3150, *n =* 220), *ens* (1.25 ± 0.28, *n =* 70), *Khc* RNAi (1.22 ± 0.44, *n =* 130). ** p<0.0l; **** p<0.0001 (Kruskal-Wallis and Dunn’s tests). For all images, scale bars: 10 µm.

**Figure S4:**
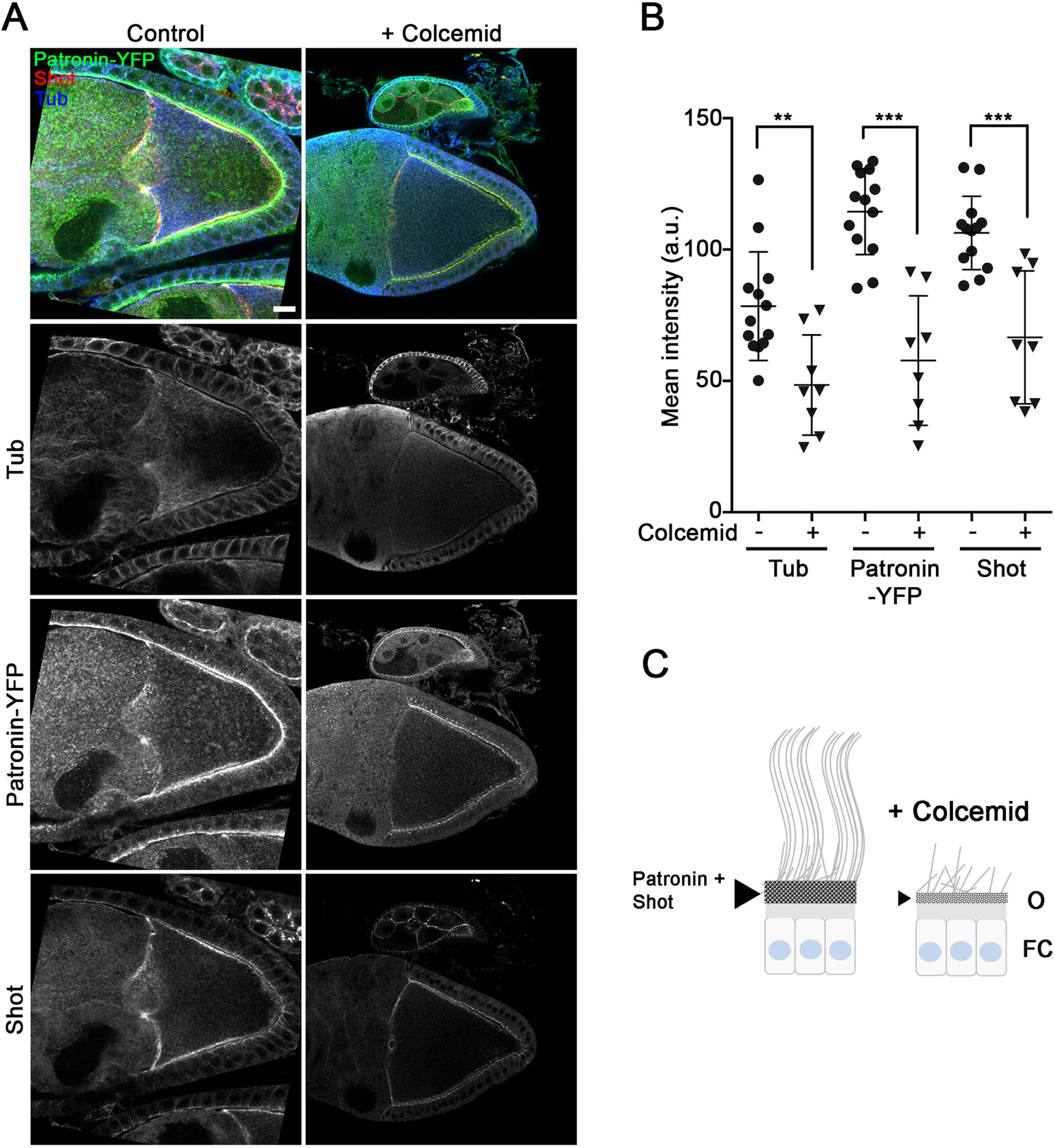
Localization of cortical ncMTOCs is MT-dependent. **(A)** Stage 9 oocytes expressing *Patronin-YFP* incubated without (Control, left panels) or with colcemid (right panel), were stained for GFP (green, and middle panels in monochrome), Shot (red, and bottom panels in monochrome) and Tub (blue and in monochrome in the second panels from the top). The colcemid treatment induces MTs depolymerization and the loss of cortical Shot and Patronin. Scale bar: 10 µm. **(B)** Dot plot showing the mean (± sd) Tubulin, Patronin-YFP and Shot signal intensities at the antero-lateral cortex, in stage 9 control or colcemid-treated oocytes. Control without colcemid (n = 13), Patronin-YFP (114.4 ± 16.32), Shot (106.3 ± 13.95) and Tub (78.41 ± 20.67); after colcemid treatment (n = 8), Patronin-YFP (57.74 ± 24.65), Shot (66.57 ± 25.31) and Tub (48.42 ± 19.09). ** p<0.0l; *** p<0.00l (Mann-Whitney test). **(C)** Scheme representing cortical ncMTOCs (Patronin + Shot) in control or colcemid-treated oocytes. O: oocyte and FC: follicle cells.

**Figure S5:**
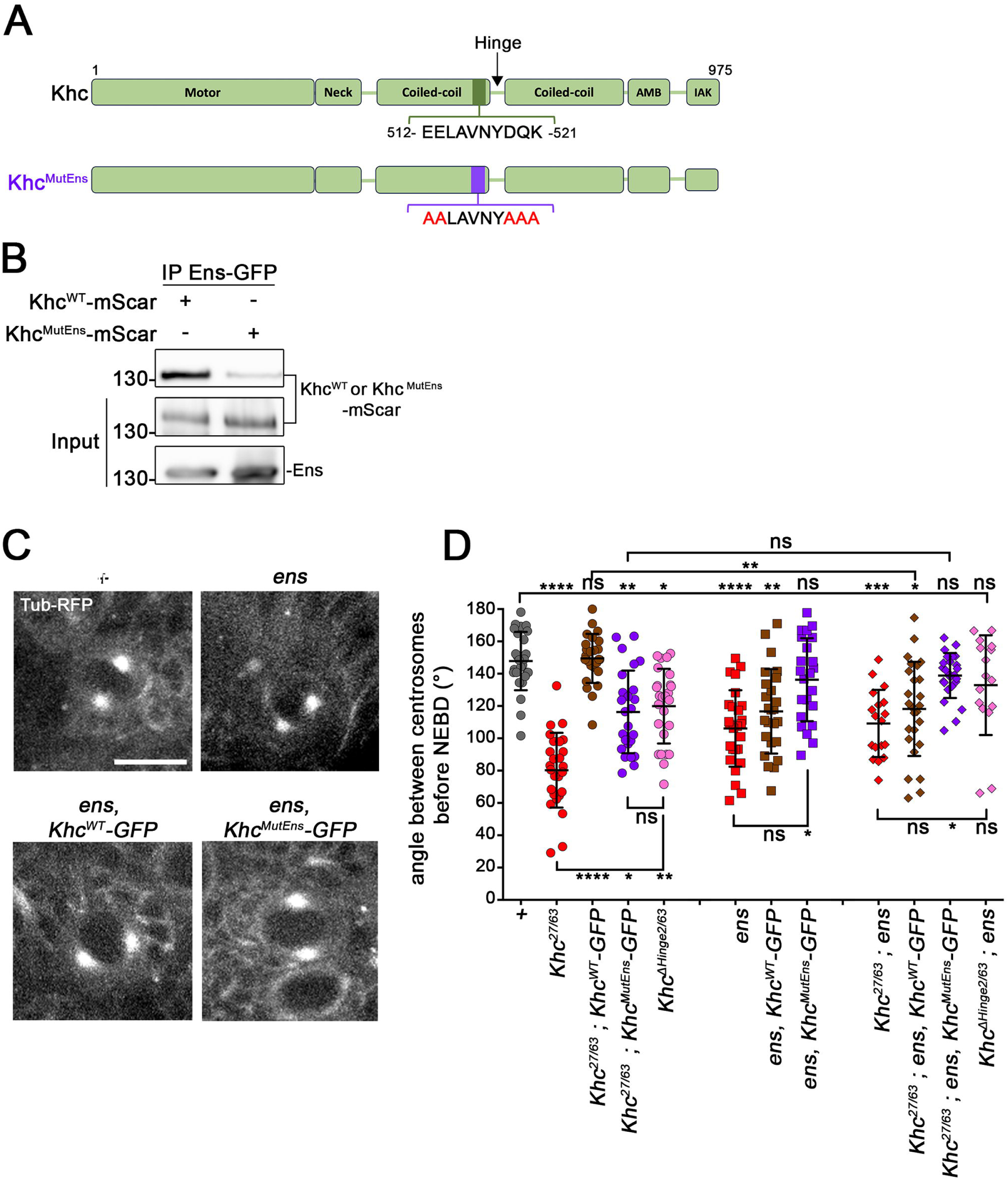
Characterization of Khc mutant defective for the interaction with Ens. **(A)** Scheme showing Khc functional domains. The motif in purple in the Khc^MutEns^ variant shows the position of the mutated Ens-binding motif in the coiled-coil 2 region. **(B)** Purified recombinant Ens^WT^-GFP proteins were immobilized on GFP-TRAP beads and tested for Khc^WT^-mScarlet or Khc^MutEns^-mScarlet binding (see Methods for details). The beads were then analyzed by Western blotting for the presence of Khc^WT^ or Khc^MutEns^. Ens^WT^-GFP binds to Khc^WT^-mScarlet and less to Khc^MutEns^-mScarlet. **(C)** Selected images showing the centrosome position 3 min before nuclear envelope breakdown (NEBD) in larval neuroblasts. The centrosomes were visualized thanks to their high microtubule nucleation activity. The neuroblasts express *Ubi-RFP-αTub* in control, *ens* and in *ens* expressing Khc^WT^-GFP or Khc^MutEns^-GFP. Scale bar: 5 µm. Images shown are 5 µm projections. **(D)** Dot plot showing the mean (± sd) centrosome separation angle in Control (147.8 ± 18.05, *n =* 26), *Khc^27/63^* (80.26 ± 23.12, *n =* 26), *Khc^27/63^* expressing Khc^WT^-GFP (149.5± 15.21, *n =* 25), *Khc^27/63^* expressing Khc^MutEns^-GFP (116.3± 25.57, *n =* 25), *Khc^ΔHinge2/63^* (119.9 ± 23.11, *n =* 26), *ens* (106.1 ± 23.64, *n =* 26), *ens* expressing Khc^WT^-GFP (116.7 ± 26.16, *n =* 25), *ens* expressing Khc^MutEns^-GFP (136.3 ± 25.66, *n* = 23), *Khc^27/63^; ens* (109.2± 20.74, *n* = 18), *Khc^27/63^;ens* expressing Khc^WT^-GFP (118.2± 29.16, *n =* 26), *Khc^27/63^; ens* expressing Khc^MutEns^-GFP (138.8± 13.92, *n =* 25), *Khc^ΔHinge2/63^; ens* (132.9± 30.84, *n =* 16). ns, not significant p>0.05; * p<0.05; ** p<0.0l; *** p<0.00l; **** p<0.000l (Kruskal-Wallis and Dunn’s tests). The centrosome separation, defective in *Khc^27/63^,* is rescued by the Khc^WT^-GFP form, and partially by Khc^MutEns^-GFP and Khc^ΔHinge2^. In an *ens* mutant background, the Khc^MutEns^-GFP rescues the centrosome separation phenotype. In a *Khc^27/63^; ens* double mutant condition, Khc^MutEns^-GFP and Khc^ΔHinge2^ rescued the centrosome separation defect.

**Video 1: Dynamics of photoconverted Dendra-Ens-decorated MTs from NC to oocyte *via* the ring canal**

Stage 9 egg chamber expressing Dendra-Ens and Pav-GFP. The time scale is shown in seconds; t = 0s corresponds to Dendra-Ens photoconversion in NCs.

Scale bar: 10 µm.

**Table S1:** Dorsal appendages phenotype and female fertility.

| Genotype | DA |  |  |  | Fertility |
| --- | --- | --- | --- | --- | --- |
|  | Normal (%) | One/Fused (%) | None (%) | n |  |
| WT | 96 | 4 | 0 | 1641 | Yes |
| GLC *Khc*<sup>27</sup> | 0 | 13 | 87 | 71 | No |
| GLC *Khc*<sup>27</sup>; *Khc*<sup>WT</sup>-GFP | 72 | 24 | 4 | 346 | Yes |
| GLC *Khc*<sup>27</sup>; *Khc*<sup>MutEns</sup>-GFP | 50 | 43 | 7 | 140 | Yes |
| *Ninein*<sup>-/-</sup> | 89 | 10 | 1 | 232 | Yes |
| *Ninein*-RFP OE | n.d. | n.d. | n.d. | n.d. | Yes |
| *ens* | 89 | 10 | 1 | 577 | No |
| *ens*<sup>WT</sup>-GFP; *ens* | 92 | 8 | 0 | 487 | Yes |
| *ens*<sup>LowMT</sup>-GFP; *ens* | 94 | 6 | 0 | 424 | Yes |
| *ens*<sup>MutKhc</sup>-GFP; *ens* | 68 | 30 | 2 | 534 | No |
| *ens*, *Khc*<sup>WT</sup>-GFP | 67 | 26 | 7 | 524 | No |
| *ens*, *Khc*<sup>MutEns</sup>-GFP | 80 | 18 | 2 | 561 | No |
| *Shot*-GFP OE; *ens* | n.d. | n.d. | n.d. | n.d. | Yes |

